# Single-Cell Multi-Omics Dissection of Malignant Evolutionary Mechanisms and Construction of a Prognostic Model for Clear Cell Renal Cell Carcinoma

**DOI:** 10.64898/2026.06.04.730125

**Authors:** Ruifei Liu, Yuchen Shi, Yuxuan Xiao, Bolin Ren, Liyuan Li, Baobin Qi, Tengyue Li, Yunpeng Zhang, Jie Gao

## Abstract

Clear cell renal cell carcinoma (ccRCC) exhibits pronounced heterogeneity across WHO histological grades, yet systematic single-cell multi-omics studies characterizing these transitions remain limited. We integrated scRNA-seq and scATAC-seq data across ccRCC WHO grades to establish a multi-omics framework encompassing tumor cells and immune populations. Using pseudotime trajectory analysis and machine learning ensembles, we developed a prognostic signature (CBG) from core nodes of transcriptional regulatory networks. We found that in tumor cells, epigenetic alterations consistently precede metabolic reprogramming and invasive adaptation. CD8^+^ T cell exhaustion followed a trajectory shifting from IRF7– to ZNF683-regulated states, while monocytes differentiated toward M1 and M2 macrophages orchestrated by NFIC/IL1B and CEBPD/GLI2. Intercellular communication networks showed a temporal progression from inflammation, through vascular remodeling, to immunosuppression dominance. The CBG signature demonstrated robust performance in independent cohorts, identifying SLC11A1 and SH3YL1 as antagonistic survival determinants. This study elucidates the dynamic molecular and immunological mechanisms underlying ccRCC grade progression, providing a robust framework for subtype-specific prognostication and precision therapeutic targeting.

## Introduction

Renal cell carcinoma (RCC) ranks among the most prevalent malignancies of the urinary system. In 2020, it accounted for 179,368 deaths worldwide^[1]^. In addition to its substantial mortality burden, the global incidence of RCC continues to rise annually^[2, 3]^. Clear cell renal cell carcinoma (ccRCC), the predominant and most aggressive histologic subtype, constitutes the majority of cases^[4, 5]^. Over one-third of ccRCC patients develop recurrence or metastasis following surgical resection, and those with metastatic disease face a dismal prognosis, with a 5-year survival rate of approximately 10%^[6]^. The WHO histological grading system (grades 1–4) remains the cornerstone of clinical prognostication in ccRCC, as grade is strongly correlated with tumor invasiveness, metastatic potential, and overall survival^[7]^. Nevertheless, conventional grading approaches reliant on histologic morphology and bulk omics profiling fail to capture the multi-pathway malignant evolution of tumor cells, transcriptional regulatory heterogeneity, and the progressive remodeling of the tumor microenvironment (TME) from a pro-inflammatory to an immunosuppressive state. This disconnect between clinical grading and underlying molecular mechanisms has hindered the advancement of precision oncology strategies^[8]^.

The advent and rapid maturation of single-cell transcriptomics (scRNA-seq) and epigenomics (scATAC-seq) have provided unprecedented single-cell resolution for dissecting tumor heterogeneity. These technologies have illuminated dynamic cellular states during tumor initiation, progression, and TME remodeling across diverse solid malignancies^[9, 10]^. In ccRCC, prior single-cell studies have predominantly focused on discrete TME compartments—such as tumor-infiltrating lymphocytes or macrophage polarization—or on features of metastatic lesions^[11]^. Systematic multi-omics integration spanning the full spectrum of WHO grades remains limited, particularly with respect to delineating tumor cell trajectory evolution, coordinated epigenetic-transcriptional regulatory networks, and the mechanistic basis of intercellular communication^[8, 12]^. These knowledge gaps impede a holistic understanding of the molecular drivers of ccRCC malignant progression and constrain the development of grade-informed prognostic models.

Prognostic assessment in ccRCC has historically depended on clinical parameters, including TNM stage, WHO grade, and Fuhrman nuclear grade, yet these yield only moderate predictive accuracy^[13]^. Recent advances in machine learning have markedly enhanced prognostic modeling in high-dimensional multi-omics datasets, enabling robust risk stratification across multiple cancer types^[14, 15]^. However, existing ccRCC prognostic signatures are largely derived from single-omics layers or conventional clinicopathologic variables, rarely incorporate core nodes from multi-perspective transcriptional regulatory networks, and lack mechanistic grounding in WHO grade-specific heterogeneity—thereby limiting model generalizability and clinical translatability^[16, 17]^.

To bridge these gaps, the present study integrates scRNA-seq and scATAC-seq datasets from the GEO repository to establish a comprehensive multi-omics framework encompassing the entire progression spectrum of ccRCC. We first systematically characterize multi-pathway malignant trajectories and stage-specific functional remodeling in tumor cells, uncovering a regulatory paradigm in which epigenetic alterations precede transcriptional activation. Second, we delineate stage-specific regulatory networks governing CD8 ⁺ T cell exhaustion differentiation and bidirectional M1/M2 polarization of macrophages. Intercellular communication analysis further elucidates the temporal transformation of the TME—from inflammation dominance, through vascular and stromal remodeling, to ultimate immunosuppression predominance. Finally, by leveraging core nodes of multi-angle transcriptional regulatory networks in conjunction with diverse machine learning ensembles, we construct and validate an optimal prognostic model (CBG) and identify key antagonistic survival markers. This work provides a systematic molecular and immunological atlas of ccRCC grade progression, establishing a robust foundation for the development of grade-specific prognostic tools and targeted therapeutic interventions.

## Methods

### Sample Information

All single-cell sequencing data used in this study were retrieved from the Gene Expression Omnibus (GEO) database^[73]^ and consisted exclusively of human samples. The dataset comprised 19 paired scRNA-seq and scATAC-seq profiles from clear cell renal cell carcinoma (ccRCC) specimens, all representing primary tumor tissues. Detailed information on accession numbers, sample types, tumor stages, and WHO histological grades is provided in Supplementary Table S1A.

Survival and prognostic modeling analyses utilized bulk RNA-seq and clinical data from The Cancer Genome Atlas (TCGA) database and the ArrayExpress database. This yielded a final cohort of 506 TCGA-KIRC patients with matched gene expression and survival information, supplemented by an independent validation set of 101 ccRCC patients with gene microarray data and corresponding clinical annotations.

### Quality Control of Single-Cell RNA Sequencing Data

Quality control of the single-cell RNA sequencing data was performed following the standard workflow of the Seurat package^[74]^ (v5.0.1). Cells were retained if they met the following criteria: (1) number of detected genes between 500 and 5,000, and (2) mitochondrial gene content <10%. After applying these filters, 112,773 high-quality cells were retained for downstream analyses.

### Preprocessing and Normalization of Single-Cell RNA Data

The standard Seurat processing pipeline was applied for dimensionality reduction and clustering. Gene expression matrices were first normalized using the “NormalizeData” function with default parameters. Highly variable features (nfeatures = 2,000) were then identified with “FindVariableFeatures”(default parameters) to select the most informative genes for subsequent analyses. Features were scaled and centered using “ScaleData”(default parameters), followed by principal component analysis (PCA) via “RunPCA”(default parameters) to capture the principal sources of variance.

### Batch Effect Correction

To mitigate potential batch effects across the 19 ccRCC samples prior to clustering, we applied the “RunHarmony” function from the Harmony package (v1.2.0) with parameters lambda = 1, max_iter = 20, and early_stop = TRUE. The Harmony-corrected embedding was used for all subsequent analyses.

### Cell Clustering and Annotation

Nonlinear dimensionality reduction was performed using “RunUMAP” on the Harmony-corrected embedding, utilizing the first 30 principal components (dims = 1:30). A shared nearest neighbor (SNN) graph was constructed with “FindNeighbors”(dims = 1:30). Clustering was conducted using “FindClusters” across a resolution range of 0.1–1.0 (resolution = seq(0.1, 1, by = 0.1)) to comprehensively evaluate cluster partitions. Clustering results were visualized and evaluated using the Clustree package (v0.5.1), and a resolution of 0.3 was selected as optimal. Marker genes for each cluster were identified with “FindAllMarkers” (only.pos = TRUE, min.pct = 0.1, logfc.threshold = 0.25). Cell types were annotated based on canonical markers from the CellMarker 2.0 database^[75]^ and established ccRCC literature^[8, 12]^. Analogous workflows were applied to characterize subpopulations within T cells and myeloid cells.

### Analysis of Single-Cell ATAC Sequencing Data

Secondary analysis of scATAC-seq data was conducted using the Signac package^[76]^, an R toolkit designed for the analysis and visualization of single-cell chromatin accessibility data. Signac integrates seamlessly with Seurat and is optimized for multimodal single-cell datasets.

To combine the 19 ccRCC samples, we first generated a consensus peak set across all samples, as independent peak calling per sample would yield inconsistent peak co-ordinates. After loading cell metadata for each sample, Fragment objects were created using the “CreateFragmentObject” function. A peak-by-cell count matrix was then generated for each sample with the “FeatureMatrix” function, and individual Seurat objects were constructed, with the corresponding Fragment objects stored in the assay. Finally, the 19 samples were merged into a single Seurat object for downstream processing.

Post-merging quality control was applied to remove low-quality nuclei. For each nucleus, we computed the fraction of fragments in peak regions, fraction of reads in genomic blacklist regions (blacklist ratio), nucleosome signal, and transcription start site (TSS) enrichment score. Following established filtering criteria^[76, 77]^, nuclei were retained if they satisfied: peak region fragments > 1,000 and < 20,000, fraction of reads in peaks > 15%, blacklist ratio < 0.05, nucleosome signal < 4, and TSS enrichment > 1. After filtering, 61,675 high-quality nuclei were retained for subsequent analyses.

Genomic coordinates for features were defined using the “GRanges” function, and gene annotations were generated based on the hg19 reference. Term frequency-inverse document frequency (TF-IDF) normalization was performed with the “RunTFIDF” function in Signac. Top features (peaks) were selected using “FindTopFeatures”, and singular value decomposition (SVD) was applied to the TF-IDF matrix via “RunSVD”. This TF-IDF + SVD workflow, also known as latent semantic indexing (LSI)^[78]^, was used for dimensionality reduction.

Harmony (v1.2.0) was then applied for batch correction (resolution = 0.5), followed by construction of a shared nearest neighbor (SNN) graph with “FindNeighbors” using dimensions 2–40. Clustering was performed with “FindClusters” at resolution = 0.5, and cluster stability was evaluated using the Clustree package. Gene activity scores were computed using the “GeneActivity” function, extending coverage to include transcription start sites and 2 kb upstream of gene bodies. Analogous workflows were applied to characterize subpopulations within T cells and myeloid cells.

### Integrated Analysis of scRNA-seq and scATAC-seq Data

Cell type annotation in scATAC-seq data is generally less definitive than in scRNA-seq due to the indirect nature of chromatin accessibility profiles. To assign cell identities in scATAC-seq, we employed two complementary strategies: (1) Label transfer within the Seurat integration framework was used to map correspondences between modalities. Anchors were identified using “FindTransferAnchors” (reduction = ‘cca’) to detect shared correlation patterns between scATAC-seq gene activity scores and scRNA-seq gene expression. Predicted cell type labels were then transferred to scATAC-seq cells via “TransferData” (weight.reduction = ‘lsi’, dims = 2:40). Cells with a maximum prediction score ≥ 0.5 were retained, yielding 32,348 high-confidence cells; (2) Cell type–specific peaks were identified from the gene activity matrix using “FindAllMarkers” in Seurat.

### Functional Reprogramming Heatmaps of Major Cell Populations

Functional enrichment analysis of the top characteristic genes/peaks from major cell populations across datasets was performed using the ClusterGVis package (v0.1.1). The top three enriched Biological Process (BP) terms were visualized (type = “BP”). Subpopulation-specific markers (genes and peaks) were identified with “FindAllMarkers” (min.pct = 0.1, logfc.threshold = 0.25).

### Copy Number Variation Analysis to Identify Malignant Tumor Cells

Copy number variations (CNVs) were inferred from scRNA-seq data using infercnv^[79]^ (v1.16.0) to distinguish malignant from non-malignant epithelial cells in ccRCC, an epithelial-derived malignancy. B cells served as the reference population (cutoff = 0.1, cluster_by_groups = TRUE, HMM = TRUE, leiden_resolution = 0.0001, denoise = TRUE). The smoothed CNV expression matrix was extracted, restricted to reference B cells and epithelial cells, and filtered to retain only genes present in both the expression matrix and a pre-prepared genomic location file.

To assess CNV heterogeneity within epithelial cells, the filtered matrix was transposed, and unsupervised k-means clustering was performed using the “kmeans” function in the stats package (v4.3.1), partitioning cells into 7 clusters^[80]^. Based on CNV heatmap visualization and overall CNV signal intensity per cluster, the cluster with the lowest CNV score was designated as low-malignancy epithelial cells and excluded from further analysis. The remaining clusters were defined as malignant tumor epithelial subpopulations. Malignant cell annotations from scRNA-seq were then propagated to scATAC-seq data using label transfer.

### Analysis of Pseudotime Trajectories of Cells

Pseudotime trajectories for malignant tumor cells, T cells, and myeloid lineages in ccRCC were constructed using Monocle3^[81]^ (v1.3.4). To address substantial batch effects among cell clusters, batch correction was applied using “align_cds” (alignment_group = “orig.ident”). CellDataSet objects were generated for tumor cells, T cells, and myeloid cells via “as.cell_data_set” from the SeuratWrappers package (v0.3.5). Dimensionality reduction was performed with “reduce_dimension” (default parameters), trajectory graphs were learned using “learn_graph” (default parameters), root nodes were manually selected, and pseudotime values were assigned with “order_cells” (default parameters). Trajectory-associated differentially expressed genes and peaks were identified using “graph_test” (neighbor_graph = “principal_graph”, cores = 8).

### Enrichment Analysis of Differentially Expressed Genes and Peaks

For differentially expressed genes identified from scRNA-seq data, functional enrichment analysis was performed using the “enrichGO” function in the clusterProfiler package^[82]^ (v4.8.3). For differentially accessible peaks from scATAC-seq, peak coordinates were converted to GRanges objects using “StringToGRanges” in Signac (v1.12.9004), followed by functional enrichment with the rGREAT package (v2.2.0) using the “great” function (Genesets = “GO:BP”, txdb = “hg19”).

### Identification of Stage-Specific Transcription Factors

Transcription factor binding motifs enriched in differentially open chromatin regions were identified using the “FindMotifs” function in Signac (default parameters). Motif enrichment was assessed by comparing the observed number of features containing each motif against the background frequency across all features, with significance determined by hypergeometric testing.

### Cis-Regulatory Element Analysis

Following previously established methodology^[8]^, co-accessibility between paired peaks was quantified using the Cicero package^[83]^ (v1.3.4.11) via graphical LASSO. The “run_cicero” function (default parameters, co-accessibility cutoff = 0.2) was used to compute Pearson correlations for peak-to-linkage pairs. Links were retained if one peak overlapped a promoter region (≤1 kb from the transcription start site), and the Pearson correlation between the average chromatin accessibility of the linked peak and the average RNA expression of the associated gene across cell types was calculated. Significant gene-linked cis-regulatory elements (cCREs) were defined as those with Benjamini–Hochberg adjusted P < 0.05 and considered candidate cCREs.

### Construction of Transcription Factor Regulatory Networks

scATAC-seq and scRNA-seq data were integrated within each cell type to identify candidate TF target genes. For each transcription factor, the top 20 target genes were defined based on two criteria: (1) direct presence of the TF binding motif in the target gene promoter, and (2) linkage of the promoter to the TF-bound peak via gene-linked cCREs.

To explore potential relationships between stage-specific differential peaks and genes, significantly dynamic genes along pseudotime (Moran’s I > 0.25) were screened. Protein–protein interaction (PPI) networks were constructed for these genes and identified target genes using the STRING database^[84]^ (v12.0) with parameters: organism = Homo sapiens, network type = full STRING network, edge meaning = evidence, minimum interaction score = 0.65. Interaction edges were overlaid onto the existing TF-target network.

Stage-specific TF regulatory networks along pseudotime were constructed using Cytoscape^[85]^ (v1.2.6) in R based on these TF-target gene pairs. Node attributes and edge representations are detailed in the figure legends.

### Cell Interaction Analysis

Intercellular communication was evaluated using MultiNicheNet^[70]^ (v2.1.0) based on ligand, receptor, and pathway information. Each condition was compared against the other two to identify condition-specific cell–cell interactions following the default MultiNicheNet workflow. Only cell clusters with ≥10 cells in at least two samples per condition were included. Top 15 ligand–receptor pairs per condition were selected for visualization in circos plots. Ligand activity inference, based on receiver cell target gene activity, was performed to evaluate potentially activated ligand signals per the default pipeline.

### Machine Learning for Prognostic Model Construction and Validation

Univariate Cox regression was performed on all candidate input genes using the “ML.Dev.Prog.Sig” function from the Mime package^[15]^ (v0.0.0.9000). Genes with P < 0.05 were retained as prognostically significant and passed to the machine learning framework.

The TCGA-KIRC dataset was randomly split into training (70%) and internal validation (30%) sets, with the E-MTAB-1980 dataset serving as the external validation cohort. The framework integrated ten classic survival algorithms: Random Survival Forest (RSF), Elastic Net (Enet), Stepwise Cox (StepCox), CoxBoost, Cox Partial Least Squares Regression (plsRcox), Supervised Principal Component (superpc), Generalized Boosted Regression Model (GBM), Survival Support Vector Machine (survivalsvm), Ridge, and Lasso. Variable selection filters included Lasso, StepCox, CoxBoost, and RSF (each with distinct hyperparameters), yielding 117 model combinations. Models were trained via K-fold cross-validation on the training set.

The model with the highest C-index in the validation set was selected as optimal due to its superior accuracy and minimized overfitting risk. Comprehensive evaluation included survival analysis across cohorts, univariate meta-analysis of 1-, 3-, and 5-year survival rates, time-dependent ROC analysis (C-index and AUC), comparison with published models(Table S2), and assessment of associations with clinical features in KIRC patients using the chi-square test.

### Survival Analysis, Correlation Analysis, and Statistical Analysis

Survival analysis for individual genes was performed using the “core_feature_sur” function from the Mime package (default parameters), which is built upon the survminer (v0.4.9) and survival (v3.3.1) packages. Patients were stratified into high– and low-expression groups based on the median gene expression level. Hazard ratios (HR), log-rank P values, and Kaplan–Meier survival curves (with 95% confidence intervals) were calculated and visualized.

Correlation between pairs of genes across different cohorts was assessed and visualized using the “cor_plot” function from the Mime package (default parameters). Pearson correlation coefficients were computed to quantify the association between variables.

## Result

### Result 1 Single-Cell Multi-Omics Decodes Heterogeneity Landscape and Core Cellular Functional Reprogramming Across WHO Grades in ccRCC

In this study, we obtained single-cell RNA sequencing (scRNA-seq) and single-cell ATAC sequencing (scATAC-seq) data from 19 paired clear cell renal cell carcinoma (ccRCC) samples (sample details provided in Supplementary Table S1). Following quality control with Seurat, 112,773 high-quality cells were retained from the scRNA-seq dataset; similarly, Signac-based quality control of scATAC-seq data yielded 61,675 cells for downstream analysis. Batch effects were corrected using Harmony^[18]^, after which all cells underwent dimensionality reduction and clustering. Cell types were annotated using the Cellmarker 2.0 database, identifying seven major populations in the scRNA-seq data: epithelial cells (CA9, CA9, CLDN3, CLDN4, FGB), NK cells (KLRD1, GNLY, FGFBP2, TRDC, KLRF1), myeloid cells (FPR1, CSF3R, LILRB2, S100A9, LILRA2), T cells (GZMK, CD27, SIT1, TNFRSF9, TRAT1), endothelial cells (ADGRL4, VWF, CLEC14A, SLCO2A1, DDL4), fibroblasts (RGS5, TAGLN, FAM162B, AGTR1, LRRC10B), and B cells (JCHAIN, IGLC2, IGHG1, IGLC3, IGHA2) (Figure 1C).

**Fig. 1.**
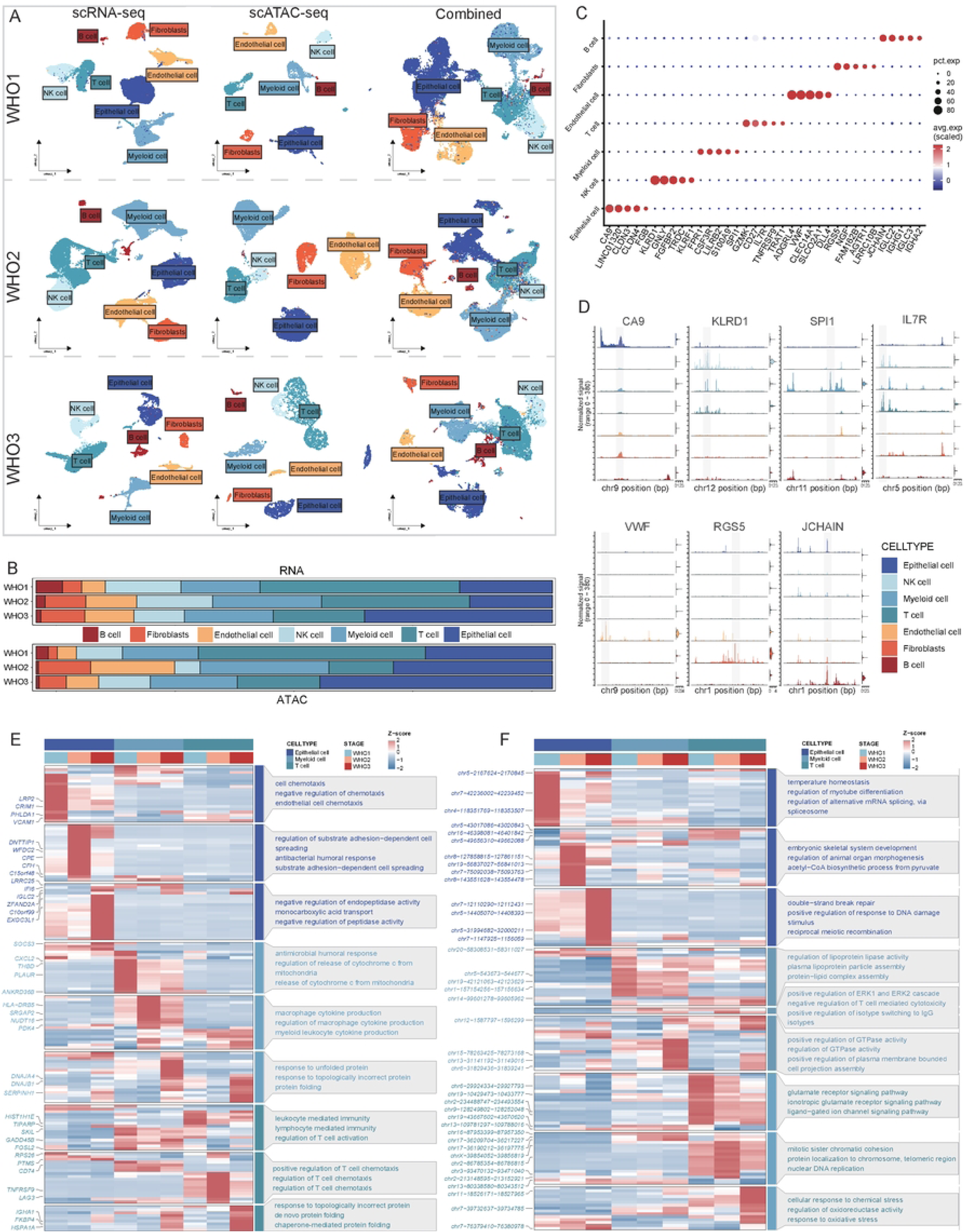
A single-cell multi-omic atlas of human clear cell renal cell carcinoma. A UMAP projections of scRNA-seq and scATAC-seq data for a total of 112,773 single cells derived from 19 ccRCC samples, stratified by global distribution and different tumor stages. B Bar plots illustrating the proportions of all cell types in scRNA-seq and scATAC-seq across different WHO grades. C Dot plot showing gene expression of cell-type marker genes in scRNA-seq data. D Normalized chromatin accessibility profiles of each cell type at canonical marker genes, with violin plots on the right indicating gene expression activity. E Signature genes of major cell types at different stages and the top 3 functional enrichment results in scRNA-seq. F Signature peaks of major cell types at different stages and the top 3 functional enrichment results in scATAC-seq. **Alt text:** Graphs and data on single-cell multi-omic analysis of human ccRCC cells, with subfigures labelled from A to F, illustrating UMAP projections, cell proportions across WHO grades, marker gene expression, chromatin accessibility profiles, and stage-specific functional enrichment results.

For the scATAC-seq dataset, we applied Seurat’s label transfer approach to assign predicted cell-type scores and retained 32,348 cells with a confidence score ≥ 0.5. Cells exhibiting substantial mixing during label transfer were classified as an “unknow” cluster and excluded from subsequent analyses.

Following annotation of both datasets, we generated single-cell UMAP projections stratified by WHO grade for scRNA-seq and scATAC-seq independently (Figure 1A). Analysis of cell-type composition across grades revealed consistent patterns in both modalities: epithelial cells, T cells, and myeloid cells constituted the three most abundant populations across all grades (Figure 1B). This concordant distribution across datasets confirms the epithelial origin of ccRCC and highlights a T cell– and myeloid cell–dominated immune microenvironment characteristic of this malignancy. We next integrated the scRNA-seq and scATAC-seq datasets and projected them into a shared low-dimensional space, demonstrating strong co-localization of corresponding cell types annotated independently in each modality (Figure 1A). Additionally, inspection of chromatin accessibility at known marker genes for each cell type further corroborated the fidelity of our annotations (Figure 1D).

To elucidate functional differences among major cell populations across ccRCC grades, we performed functional enrichment analysis on marker peaks identified in scATAC-seq and marker genes detected in scRNA-seq (Figure 1E,F).

Functional enrichment analysis revealed pronounced reprogramming across the three major cell populations in ccRCC as a function of WHO histological grade. In epithelial cells, a characteristic “homeostasis-to-stress” phenotypic shift was observed. Multi-omics integration of epithelial cells showed that chromatin accessibility in WHO grade 1 was predominantly enriched in pathways related to mRNA alternative splicing regulation, whereas the transcriptome was enriched in cell chemotaxis-associated processes, consistent with maintenance of epithelial homeostasis at early stages. In WHO grade 2, open chromatin regions centered on metabolic activation pathways, while the transcriptome highlighted cell adhesion and immune-related functions, indicative of structural remodeling and enhanced functional activity. By WHO grade 3, scATAC-seq prominently featured DNA damage repair pathways, with the transcriptome enriched in metabolic regulation and proteolysis-related processes, suggesting adaptive responses to stressful microenvironmental conditions.

In myeloid cells, WHO grade 1 exhibited chromatin accessibility enriched in ATP metabolic processes, purine ribonucleoside triphosphate metabolic processes, and membrane fission—pathways linked to energy supply and membrane dynamics. These epigenetic changes supported transcriptome enrichment in chemokine-mediated signaling and granulocyte chemotaxis/migration, defining an early-stage signature of “energy activation–chemotaxis recruitment.” At WHO grade 2, open chromatin focused on regulation of the ERK1/2 cascade and negative regulation of T cell cytotoxicity, driving transcriptome-level production of macrophage-associated cytokines and reflecting features of mid-stage immune crosstalk and competition. In WHO grade 3, scATAC-seq highlighted regulation of GTPase activity, which corresponded to transcriptome enrichment in unfolded protein response and related functions, suggesting that late-stage myeloid cells maintain protein homeostasis through coordinated epigenetic-transcriptional mechanisms while responding to microenvironmental stress.

T cells displayed pronounced reprogramming toward immune functionality across WHO grades. In WHO grade 1, chromatin accessibility was enriched in the glutamate receptor signaling pathway, while the transcriptome was dominated by processes related to T cell activation regulation and broader immune response functions, consistent with preservation of an active immune state during early disease stages. In WHO grade 2, open chromatin regions centered on nuclear DNA replication pathways, which corresponded to transcriptome enrichment in T cell chemotaxis regulation, indicative of a mid-stage functional shift toward microenvironmental adaptation. By WHO grade 3, scATAC-seq prominently featured the oxidative stress response pathway, aligned with transcriptome enrichment in the unfolded protein response, suggesting that late-stage T cells experience heightened stress and transition into a state of persistent exhaustion.

### Result 2 Multi-Omics Dissection of Multi-Path Malignant Evolutionary Trajectories in ccRCC Tumor Cells Reveals Stage-Specific Functional Remodeling and Transcriptional Regulatory Networks

During malignant progression of clear cell renal cell carcinoma (ccRCC), tumor cells display complex differentiation states and dynamic transformation trajectories. We first assessed copy number variations across all epithelial cells in the scRNA-seq dataset using inferCNV and classified cells exhibiting significant copy number aberrations as malignant tumor cells. Downstream analyses were then restricted to this malignant population(Supplementary Figure S1A). Subsequently, we applied Seurat’s label transfer approach to propagate tumor cell annotations from the scRNA-seq to the scATAC-seq dataset.

We next constructed differentiation trajectories for tumor cells independently in the scRNA-seq and scATAC-seq datasets. Using Monocle3, tumor cells were reclustered, and their branching nodes were tracked (Figure 2A, B). Notably, both datasets revealed a shared major trajectory corresponding to progressive differentiation from WHO grade 1 to grade 2 and ultimately to grade 3 (RNA-Fate1 and ATAC-Fate1). However, dataset-specific trajectories were also identified: transcriptomic analysis delineated an evolutionary path from WHO grade 2 to grade 3 (RNA-Fate2), whereas epigenomic analysis captured a transition from WHO grade 1 to grade 2 (ATAC-Fate3). These findings indicate dynamic grade-dependent changes in tumor cells across both modalities, yet the distinct trajectory patterns suggest differential regulatory mechanisms operating at the transcriptional and epigenetic levels.

**Fig. 2.**
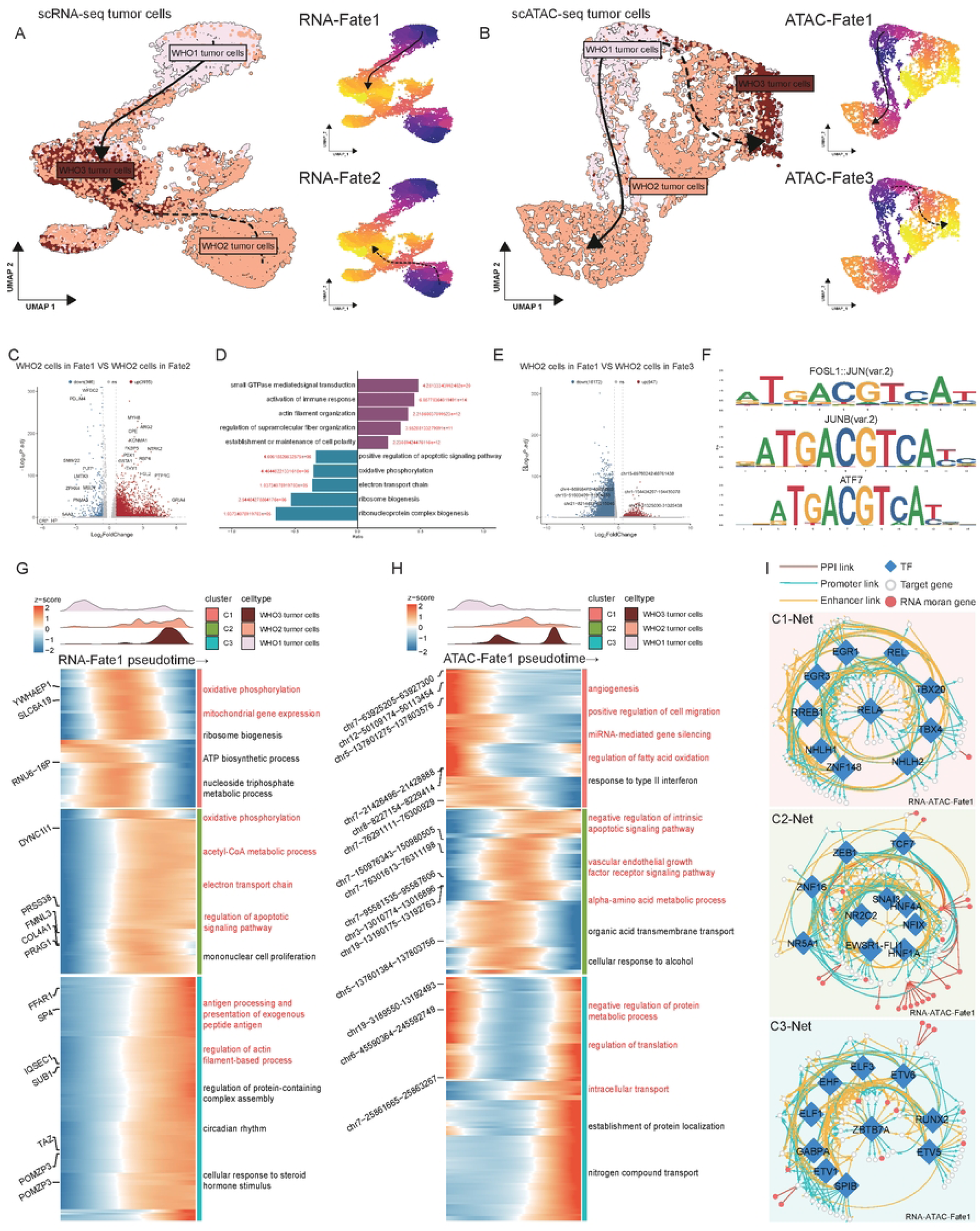
Grade-specific regulatory elements and chromatin accessibility landscapes in ccRCC tumor cells. A UMAP projection (left) and pseudotime trajectories (right) of tumor cells in scRNA-seq data. Cells are colored by WHO stage, with dark and light colors representing the origin and endpoint of differentiation, respectively. Arrows indicate the direction of differentiation. B UMAP projection (left) and pseudotime trajectories of distinct differentiation paths (right) of tumor cells in scATAC-seq data. C Volcano plot of differentially expressed genes between WHO stage 2 cells in Fate1 and Fate2 trajectories from scRNA-seq. Significant genes are determined by log2_fc > 0.25. D GO enrichment analysis of downregulated (left) and upregulated (right) genes identified from the volcano plot in C. E Volcano plot of differentially accessible peaks between WHO stage 2 cells in Fate1 and Fate3 trajectories from scATAC-seq, with a threshold of log2_fc > 0.25. F Transcription factor (TF) motif enrichment analysis of upregulated peaks identified from the volcano plot in E. G Combined heatmap revealing genes/peaks with significantly altered expression during tumor cell progression in scRNA-seq. The top panel shows the cell density cloud plot, and the right panel presents the GO terms enriched in different clusters. H Combined heatmap revealing peaks/genes with significantly altered accessibility during tumor cell progression in scATAC-seq. The top panel shows the cell density cloud plot, and the right panel presents the GO terms enriched in different clusters. I Regulatory network in clusters C1, C2, and C3. Arrows represent distinct regulatory modes. Different shapes represent gene identities: diamonds represent transcription factors, white circles represent target genes, and red circles represent genes with Moran’s I > 0.25 from Figure 2G. **Alt text:** Graphs and data on grade-specific regulatory elements and chromatin accessibility in ccRCC, with subfigures labelled from A to I, illustrating pseudotime trajectories (Fate1/2/3), volcano plots of DEGs and DAPs, TF motif enrichment, progression-related heatmaps, and a multi-omic regulatory network.

We first examined differences among tumor cells of the same grade along divergent differentiation paths. Along RNA-Fate1, WHO grade 2 cells upregulated genes enriched in small GTPase-mediated signal transduction, actin filament organization, and establishment or maintenance of cell polarity—features consistent with an earlier or intermediate differentiation state characterized by heightened signaling sensitivity, plasticity, and migratory potential. In contrast, along RNA-Fate2, these cells showed enrichment in metabolic and protein synthesis pathways, suggesting completion of partial differentiation and entry into a stage of elevated metabolic activity and functional terminalization (Figure 2C, D).

To investigate epigenetic differences underlying divergent trajectories, we compared chromatin accessibility in WHO grade 2 tumor cells along ATAC-Fate1 versus ATAC-Fate3 in the scATAC-seq dataset. Peaks upregulated in ATAC-Fate1 WHO grade 2 cells (relative to those stalled in ATAC-Fate3) were significantly enriched for motifs of FOSL1::JUN, JUNB, and ATF7—all members of the AP-1 transcription factor family. These factors are well-established regulators of stress responses, cell proliferation, and tumor-associated signaling pathways, suggesting that AP-1 signaling may play a critical role in driving the malignant progression of tumor cells from WHO grade 2 to grade 3 through these functional axes (Figure 2E, F).

Taken together, at the transcriptional level, tumor cells along the RNA-Fate1 trajectory retained signal activation and immune-related plasticity at WHO grade 2, whereas those along RNA-Fate2 exhibited features of elevated metabolic synthesis and functional maturation. At the epigenetic level, tumor cells following the ATAC-Fate1 trajectory were driven by AP-1 family motifs to regulate stress responses and proliferation, thereby promoting malignant advancement, while those along ATAC-Fate3 remained relatively arrested in an early-to-mid differentiation state.

To investigate functional remodeling along the complete differentiation trajectory of tumor cells, we performed pseudotime-based functional enrichment analyses on RNA-Fate1 and ATAC-Fate1. Differentially expressed genes (Moran’s I > 0.05) were clustered into three groups—C1, C2, and C3—corresponding to hierarchical stages of cellular progression. Pseudotime density plots for each cell population were overlaid to associate specific functional programs with these clusters (Figure 2G, H).

Notably, scATAC-seq data exhibited a greater propensity to prospectively reveal the “blueprint” or preparatory epigenetic states underlying malignant progression in tumor cells. In the C1 stage of tumor evolution, while scRNA-seq primarily captured intrinsic high metabolic activity—such as oxidative phosphorylation and mitochondrial gene expression—to sustain rapid proliferation, scATAC-seq identified open chromatin regions poised for future microenvironmental interactions and invasiveness. These included regulatory elements linked to angiogenesis, cell migration, miR-NA-mediated gene silencing, and lipid metabolism. This pattern indicates that epigenetic reprogramming precedes detectable transcriptional changes, establishing a pre-adaptive landscape that primes tumor cells for subsequent phenotypic transitions.

As progression advanced to the C2 stage, this epigenetic “preparation” exhibited stronger coupling with transcriptional alterations. Open chromatin sites identified by scATAC-seq in the VEGF signaling pathway, apoptosis inhibition, and amino acid/organic acid metabolism aligned closely with transcriptome features detected by scRNA-seq, including oxidative phosphorylation, acetyl-CoA metabolism, electron transport chain activation, and regulation of apoptosis signaling pathways. These findings suggest that the mid-stage represents a phase of tight epigenetic-transcriptional coordination, in which epigenetic regulatory elements directly orchestrate gene expression programs that support cell survival, metabolic reprogramming, and angiogenesis.

In the C3 stage, chromatin accessibility changes detected by scATAC-seq were predominantly enriched in pathways governing protein metabolism regulation, translation, and intracellular transport—processes associated with largescale cellular structural and functional reconfiguration. In parallel, scRNA-seq more directly captured the functional consequences of these changes, including antigen processing and presentation (linked to immune evasion) and regulation of actin filament-based processes (associated with cytoskeletal rearrangement, invasion, and metastasis), with marked upregulation of relevant genes. Collectively, these results indicate that late-stage tumor cells achieve acquisition of key malignant phenotypes at the transcriptional and protein levels through epigenetic-driven functional remodeling.

Overall, functional enrichment analyses from scRNA-seq and scATAC-seq were highly concordant, delineating a broad evolutionary trajectory in tumor cells from metabolic reprogramming to the acquisition of immune evasion and invasive phenotypes. Together, these modalities outlined a dynamic process whereby tumor cells progressively adapt to and actively reshape their microenvironment.

To elucidate the core regulatory mechanisms underpinning this staged functional evolution, we constructed stage-specific transcriptional regulatory networks based on differential chromatin accessibility, differentially expressed genes, and transcription factor target relationships at each phase. Each network exhibited distinct central nodes that closely aligned with the functional attributes of the corresponding stage, collectively forming a temporal regulatory axis driving malignant tumor cell evolution (Figure 2I).

In the early-stage network (C1-Net), RELA (also known as p65) emerged as the central hub. As a critical subunit of the canonical NF-KB pathway, RELA is constitutively activated in ccRCC and has been shown in multiple studies to promote inflammatory responses, cell survival, metabolic reprogramming, and immune evasion through regulation of downstream targets[19–21]. In the mid-stage network (C2-Net), SLC6A3 occupied the central position as a target gene regulated by multiple transcription factors. SLC6A3 encodes a dopamine transporter^[22, 23]^, is associated with HIF pathway activation, and serves as one of the specific biomarkers of ccRCC. In the late-stage network (C3-Net), ZBTB7A constituted the core node. This factor has been established as an oncogene across various malignancies, where it recruits repressor complexes to suppress cell-cycle inhibitors (p21 and p53), thereby promoting proliferation and survival. In renal cancer, ZBTB7A is highly expressed, drives tumor cell proliferation and invasion, and establishes a positive feedback loop^[24, 25]^.

In summary, integrative analysis of scRNA-seq and scATAC-seq uncovered multiple parallel differentiation trajectories in ccRCC tumor cells across WHO grade progression. The transcriptome revealed a gradient from early signal activation and immune plasticity to late-stage elevated metabolism and functional maturation, whereas the epigenome highlighted progression from early metabolic reprogramming to AP-1-driven stress and proliferative responses. Pseudotime analyses demonstrated that epigenetic alterations frequently precede and establish the foundation for subsequent malignant phenotypes. Stage-specific regulatory networks identified RELA, SLC6A3, and ZBTB7A as pivotal core nodes, collectively orchestrating a temporal regulatory axis governing tumor cell malignant evolution.

### Result 3 Multi-Omics Unveils Stage-Specific Regulatory Networks and Immune Checkpoint Co-Expression Mechanisms Driving CD8⁺T Cell Exhaustion Differentiation Trajectories in ccRCC

Given the marked heterogeneity and functional state differences observed in tumor cells across divergent evolutionary pathways, we next sought to delineate the dynamic evolution of immune cells within the tumor microenvironment (TME), with a specific focus on T cell progression. As key effectors of antitumor immunity, CD8⁺ T cells undergo a critical transition from naive to exhausted states that can profoundly influence tumor immune evasion and disease progression. Using scRNA-seq data, we first annotated T cells into distinct subtypes based on canonical marker genes, identifying five major subpopulations: naive CD4⁺ T cells, naive CD8⁺ T cells, regulatory T cells (Treg), exhausted CD8⁺ T cells, and proliferative exhausted CD8⁺ T cells. Subsequently, by integrating label transfer with manual curation, we annotated three principal subtypes in the scATAC-seq dataset: naive CD4⁺ T cells, naive CD8⁺ T cells, and exhausted CD8⁺ T cells (Supplementary Figure S1B,C,D).

Building on these annotations, we applied the same pseudotime trajectory inference strategy to both omics datasets for T cells. Transcriptomic trajectory analysis revealed a progressive differentiation route in CD8⁺ T cells, evolving from naive CD8⁺ T cells to exhausted CD8⁺ T cells and ultimately to proliferative exhausted CD8⁺ T cells (RNA-Fate1) (Figure 3A). Epigenomic trajectory inference showed a comparable pattern, with naive CD8⁺ T cells gradually transitioning into exhausted CD8⁺ T cells along pseudotime (ATAC-Fate1) (Figure 3B). Accordingly, subsequent analyses focused on the shared CD8⁺ T cell differentiation trajectory conserved across both modalities, em-ploying the identical analytical framework as previously described.

**Fig. 3.**
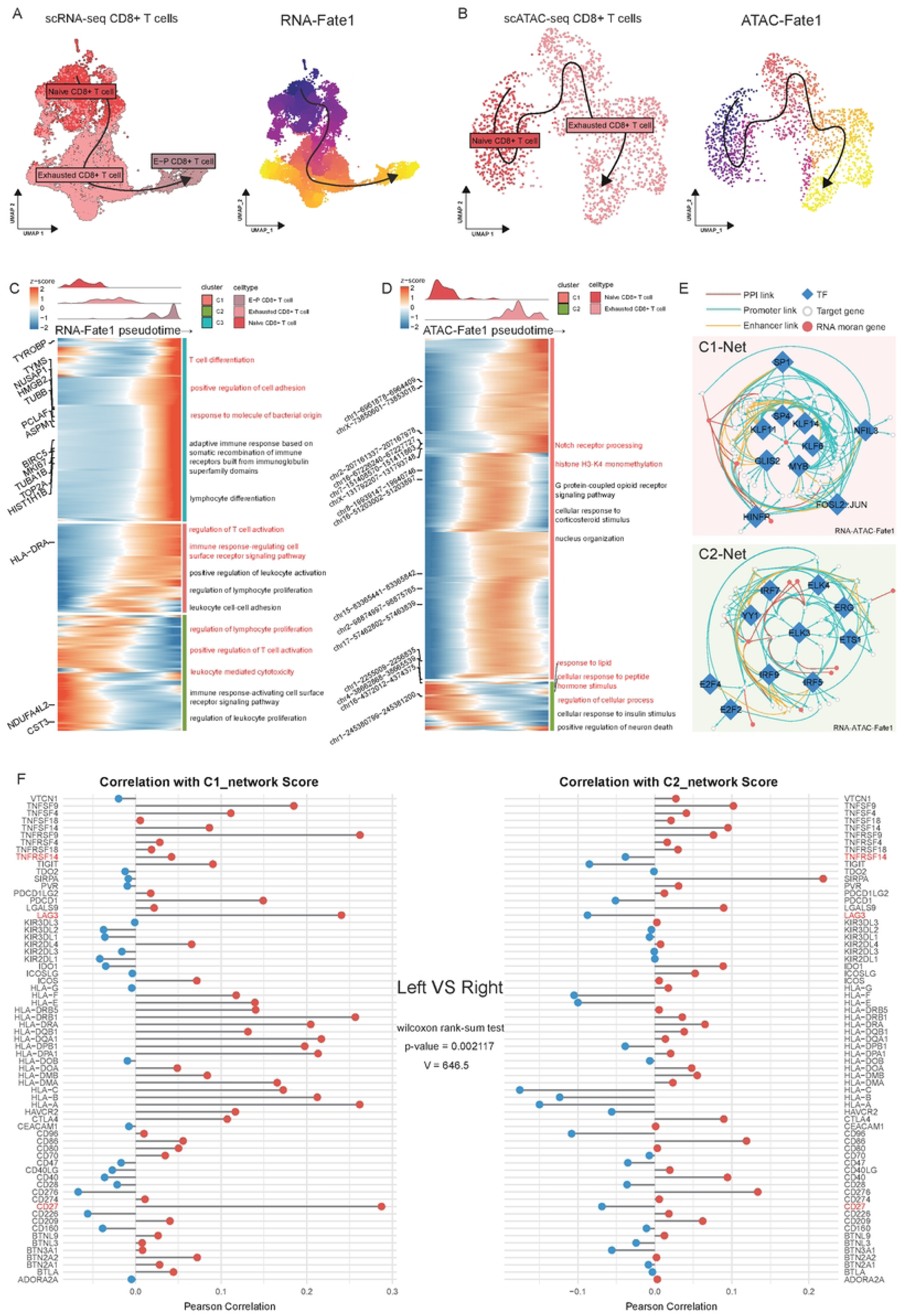
Integrative multi-omics reveals dynamic transcriptional and epigenetic regulation of T cell differentiation. A UMAP projection (left) and pseudotime trajectories (right) of CD8+ T cells in scRNA-seq and scATAC-seq data. Cells are colored by cell type, with dark and light colors representing the origin and endpoint of differentiation, respectively. Arrows indicate the direction of differentiation. B UMAP projection (left) and pseudotime trajectories of distinct differentiation paths (right) of CD8+ T cells in scATAC-seq data. C Combined heatmap revealing genes/peaks with significantly altered expression during CD8+ T cell progression in scRNA-seq. The top panel shows the cell density cloud plot, and the right panel presents the GO terms enriched in different clusters. D Combined heatmap revealing genes/peaks with significantly altered accessibility during CD8+ T cell progression in scATAC-seq. The top panel shows the cell density cloud plot, and the right panel presents the GO terms enriched in different clusters. E Regulatory network in cluster C1 and C2. Arrows represent distinct regulatory modes. Different shapes represent gene identities: diamonds represent transcription factors, white circles represent target genes, and red circles represent genes with Moran’s I > 0.25 from Figure 3C. F Pearson correlation analysis between C1-Net (left) and C2-Net (right) with the expression levels of 67 immune checkpoint genes (ICGs). **Alt text:** Graphs and data on integrative multi-omic analysis of CD8+ T cell differentiation, with subfigures labelled from A to F, illustrating pseudotime trajectories, progression-related heatmaps (RNA/ATAC), regulatory network, and correlation with ICG expression.

Functional enrichment results revealed distinct stage-specific features along the CD8⁺ T cell pseudotime trajectory. In the C1 stage, a characteristic “multi-omics functional synergy initiation” was evident. At the transcriptional level, CD8⁺ T cells were enriched in pathways governing lymphocyte proliferation regulation, positive regulation of T cell activation, and cell-mediated cytotoxicity, indicating initiation of early activation and immune recognition programs with retained proliferative and cytotoxic potential. Correspondingly, in the early epigenetic landscape, open chromatin regions were predominantly enriched in metabolic pathways such as lipid response and peptide hormone stimulation. This suggests that chromatin had already opened regulatory elements linked to metabolic sensing prior to robust transcriptional activation, thereby providing epigenetic priming to support rapid proliferation and effector function establishment. These findings imply that metabolic cues may drive the onset of early immune programs through epigenetic mechanisms, with the two layers cooperating to establish the foundational functional state of CD8⁺ T cells (Figure 3C, D).

Upon progression to the C2 stage, the dominant feature shifted to “epigenetic regulation-driven functional transformation.” Transcriptionally, CD8⁺ T cells transitioned toward regulation of T cell activation and immune response pathways mediated by cell surface receptor signaling, consistent with entry into an activation-imbalanced state characterized by inhibitory receptor upregulation and early exhaustion features, while retaining partial residual activity. In the corresponding later epigenetic profile, open chromatin was enriched in pathways involving Notch receptor processing, his-tone H3-K4 mono-methylation, and other epigenetic modifications as well as signal transduction cascades. This indicates substantial chromatin remodeling, with epigenetic regulation and signal pathway reconfiguration emerging as primary drivers. Collectively, these observations suggest that epigenetic landscape remodeling and up-stream signal modulation represent the key mechanisms precipitating the mid-stage exhaustion phenotype observed at the transcriptional level—namely, through targeted epigenetic modifications that orchestrate downstream gene expression to promote the transition of CD8⁺ T cells from a functional to an exhausted state.

The C3 stage was uniquely prominent in the transcriptome, where CD8⁺ T cells showed enrichment in pathways related to T cell differentiation, positive regulation of cell adhesion, and responses to bacterially derived molecules. This profile indicates that, despite residing in an exhausted state, this subpopulation retains proliferative capacity and corresponds to non-terminally exhausted cells, consistent with prior reports.

Integrating multi-omics profiles across these stages delineated a clear differentiation trajectory of CD8⁺ T cells from functional activation to early exhaustion. At its core lies a “regulation–function” dynamic adaptation mechanism: epigenetic alterations uncovered by scATAC-seq (chromatin opening and signal reconfiguration) and functional phenotypes captured by scRNA-seq (proliferative activation and exhaustion signatures) co-evolve and mutually reinforce throughout pseudotime progression, providing complementary evidence from the regulatory foundation and phenotypic expression levels.

To further dissect the core transcriptional mechanisms underlying the staged functional and regulatory features described above, we applied the same strategy to construct stage-specific regulatory networks. We built a transcription factor (TF) regulatory network specifically for the CD8⁺ T cell differentiation trajectory, identifying pivotal regulatory nodes at each critical stage and delineating the regulatory chain linking “epigenetic regulation – gene expression – cellular function.”

In the early-stage regulatory network (C2-Net), IRF7 emerged as the central hub. Mechanistic studies have demonstrated that the IFNγ-STAT1-IRF7 axis induces IFI35 expression, which in turn is associated with enhanced CD8⁺ T cell infiltration^[26]^. In the late-stage regulatory network (C1-Net), ZNF683 occupied the central regulatory position as a target gene modulated by multiple transcription factors. Prior research has established ZNF683 as a marker of an intermediate exhausted state within tumor-infiltrating CD8⁺ T cell subpopulations—distinct from terminal exhaustion—and linked its expression to responsiveness to PD-1 blockade therapy, corroborating our earlier observations. Moreover, pseudotime analysis of the transcriptome revealed ZNF683 as a significantly dynamic gene (Moran’s I > 0.25), underscoring its potential key role in orchestrating CD8⁺ T cell differentiation (Figure 3E).

In summary, CD8⁺ T cell differentiation is characterized by a phased transition of core regulatory nodes from IRF7 to ZNF683. IRF7 drives early immune activation and the establishment of infiltration potential, whereas ZNF683 governs late-stage exhaustion progression and plasticity maintenance. These nodes align closely with the multi-omics regulatory signatures observed at distinct stages, collectively delineating the transcriptional regulatory framework that governs the transition of CD8⁺ T cells from a functional to an exhausted state and providing precise molecular evidence for the process of T cell exhaustion.

Immune checkpoint molecules play a pivotal role in modulating T cell activation, functional maintenance, and exhaustion development. Their aberrant engagement constitutes a major molecular basis for CD8⁺ T cell dysfunction in the tumor microenvironment. Thus, examining the correlation patterns between CD8⁺ T cell regulatory networks and immune checkpoint molecules across distinct functional states is essential for understanding the mechanisms governing T cell exhaustion.

We further assessed the correlation characteristics between stage-specific regulatory networks and immune checkpoint molecules. Using a curated set of 67 canonical immune checkpoint genes reported in the literature, we calculated the distribution of correlations between these genes and the key regulatory network genes in the two states. Comparative analysis revealed that the overall correlation between C1-Net and immune checkpoint genes was significantly higher than that observed for C2-Net. Both median and mean correlation values were markedly elevated in C1-Net (Wilcoxon rank-sum test, V = 646.5, p = 0.002117). These findings indicate that, during the early functional stage, immune checkpoint molecules exhibit relatively weak association with the regulatory network, likely facilitating the maintenance of T cell activation homeostasis and cytolytic capacity. In contrast, upon entry into the exhausted state, the correlation with key network genes substantially increases, suggesting that immune checkpoint molecules may synergize with exhaustion-associated regulatory circuitry to reinforce immunosuppressive phenotypes and exacerbate T cell functional impairment.

To further delineate the integration patterns of immune checkpoint molecules within stage-specific networks, we examined the correlation profiles of overlapping genes present in both networks (Figure 3F). Notably, CD27 and TNFRSF14 were shared between C2-Net and C1-Net, yet their correlations with overall network expression exhibited completely opposing directions. In C2-Net, both molecules displayed negative correlations, suggesting that these co-stimulatory factors may be suppressed within the activation-regulatory network, thereby helping to sustain effector response homeostasis and prevent premature exhaustion entry. Consistent with prior reports, CD27, a TNFR superfamily member, promotes CD8⁺ T cell proliferation, survival, and memory formation^[27]^; its negative correlation may limit excessive activation and avert functional collapse. Similarly, TNFRSF14 enhances T cell activation and tumor infiltration via the HVEM/LTβR axis^[28]^, and its negative correlation likely reflects a balanced regulatory role during functional initiation to support initial cytotoxicity without engaging inhibitory circuits. In C1-Net, these correlations reversed to positive, indicating reintegration of these molecules into exhaustion-associated regulatory circuits, potentially preserving partial proliferative capacity and plasticity in intermediate exhausted cells and providing a molecular foundation for responsiveness to immune checkpoint blockade. Specifically, CD27 expression in exhausted CD8⁺ T cells has been shown to promote self-renewal of progenitor-like subpopulations and synergize with anti-PD-1 to reverse exhaustion^[29]^, while elevated TNFRSF14 enhances CAR-T cell function and alleviates immunosuppression in the tumor microenvironment^[30]^; the positive correlation may reinforce exhausted-cell plasticity.

In addition, TNFRSF9 exhibited positive correlations in both networks, with significantly stronger correlation strength in C1-Net compared to C2-Net. This gradient amplification pattern reflects the progressive and increasingly tight integration of TNFRSF9 with the regulatory network during exhaustion progression, consistent with its established role in promoting proliferation of Tex precursor cells and driving transition toward terminal exhaustion. Literature indicates that TNFRSF9 independently drives proliferation and terminal differentiation of exhausted CD8⁺ T cells via the NF-KB pathway^[31]^; its stronger late-stage correlation may contribute to stabilization of the exhaustion phenotype.

These differential correlation patterns underscore the dynamic remodeling of checkpoint molecules along the CD8⁺ T cell differentiation trajectory: certain molecules (e.g., CD27 and TNFRSF14) undergo directional switching that may drive the functional transition from effector to exhausted states, while others (e.g., TNFRSF9) display amplified late-stage correlations, further confirming stronger functional linkage between checkpoint genes and the regulatory network during exhaustion.

By integrating multi-omics pseudotime trajectories from scRNA-seq and scAT-AC-seq, we delineated the differentiation dynamics of CD8⁺ T cells in the tumor microenvironment from naive to exhausted states: C1 features multi-omics synergistic initiation of functional activation; C2 undergoes epigenetic remodeling that drives transition to early exhaustion; C3 retains proliferative potential characteristic of non-terminal exhaustion. The core transcriptional regulatory node shifts from IRF7 to ZNF683, forming a sequential epigenetic-transcriptional-functional chain. Immune checkpoint analysis further reveals significantly higher correlation of checkpoint genes in C1-Net than in C2-Net, with CD27 and TNFRSF14 showing negative correlations in the early stage that reverse to positive in the later stage, and TNFRSF9 exhibiting a gradient of enhanced correlation in the late stage. These dynamic patterns highlight the heterogeneity of CD8⁺ T cell exhaustion and the “pull-push” balance mechanism, offering molecular insights for precision immunotherapy.

### Result 4 Multi-Omics Elucidates Functional Polarization Features and Stage-Specific Transcriptional Regulatory Networks in M1/M2 Biphasic Differentiation Trajectories of Macrophages in ccRCC

Myeloid cells are pivotal regulators within the tumor microenvironment and undergo dynamic phenotypic shifts during the immune response. To elucidate these changes, we analyzed the evolutionary trajectories of myeloid cell subpopulations. Using scRNA-seq data, we annotated myeloid cells into six major subtypes: monocytes, dendritic cells, mast cells, M1 macrophages, M1/M2 mixed-type macrophages, and M2 macrophages. Subsequently, integrating label transfer with manual curation, we identified four principal subtypes in the scATAC-seq dataset: monocytes, dendritic cells, M1 macrophages, and M2 macrophages (Supplementary Figure S1E,F,G).

Macrophage functional polarization represents the directed differentiation toward proinflammatory (M1) or immunosuppressive (M2) phenotypes in response to tumor microenvironmental cues. In the tumor immune milieu, M1 macrophages typically exert proinflammatory effects and enhance antitumor immunity through cytokines such as TNF-α and IL-1β, whereas M2 macrophages promote tumor progression and immune evasion via immunosuppressive mechanisms. Given that M1/M2 mixed-type cells lacked a clear unidirectional trajectory signature and that dendritic cells and mast cells do not derive from the monocyte lineage, these three populations were excluded from subsequent pseudotime analyses. Focusing on monocytes as the origin, we examined the divergent differentiation paths toward M1 macrophages (RNA/ATAC-Fate1) and M2 macrophages (RNA/ATAC-Fate2) (Figure 4A, B).

**Fig. 4.**
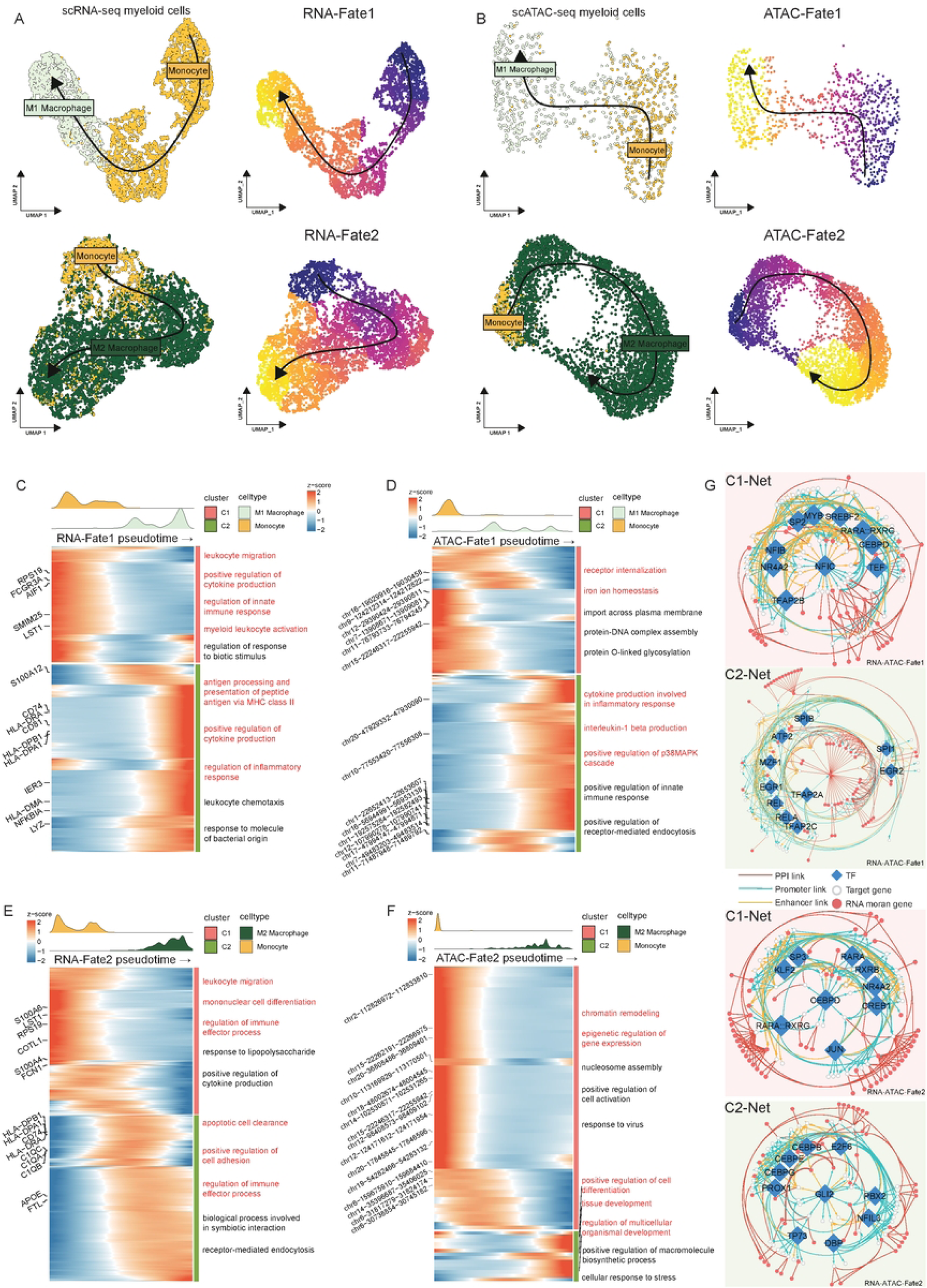
Co-evolutionary transcriptomic and epigenetic map of monocyte-to-macrophage differentiation. A UMAP projection (left) and pseudotime trajectories (right) of monocytes and macrophages in scRNA-seq and scATAC-seq data. Cells are colored by cell type, with dark and light colors representing the origin and endpoint of differentiation, respectively. Arrows indicate the direction of differentiation. B UMAP projection (left) and pseudotime trajectories of distinct differentiation paths (right) of monocytes and macrophages in scATAC-seq data. C Combined heatmap revealing genes/peaks with significantly altered expression during monocyte and M1 macrophage progression in scRNA-seq. The top panel shows the cell density cloud plot, and the right panel presents the GO terms enriched in different clusters. D Combined heatmap revealing genes/peaks with significantly altered accessibility during monocyte and M1 macrophage progression in scATAC-seq. The top panel shows the cell density cloud plot, and the right panel presents the GO terms enriched in different clusters. E Combined heatmap revealing genes/peaks with significantly altered expression during monocyte and M2 macrophage progression in scRNA-seq. The top panel shows the cell density cloud plot, and the right panel presents the GO terms enriched in different clusters. F Combined heatmap revealing genes/peaks with significantly altered accessibility during monocyte and M2 macrophage progression in scATAC-seq. The top panel shows the cell density cloud plot, and the right panel presents the GO terms enriched in different clusters. G Regulatory network in clusters C1 and C2. Arrows represent distinct regulatory modes. Different shapes represent gene identities: diamonds represent transcription factors, white circles represent target genes, and red circles represent genes with Moran’s I > 0.25 from Figure 4C and 4E. The upper and lower parts display the networks for Fate1 and Fate2, respectively. **Alt text:** Graphs and data on monocyte-to-macrophage differentiation, with subfigures labelled from A to G, illustrating pseudotime trajectories (M1/M2 fates), progression-related heatmaps for both transcriptomic and epigenetic layers, and fate-specific regulatory networks.

Multi-omics analysis revealed stage-specific co-features across these two core trajectories. In the early phase, RNA-Fate1 and RNA-Fate2 exhibited concordant activity, both enriched in immune-related preparatory functions, including leukocyte migration and regulation of immune effector processes—indicative of shared transcriptional priming for immune cell recruitment and effector activation. At the epigenetic level, however, trajectory-specific pre-adaptation was apparent: ATAC-Fate1 featured open regulatory elements associated with internalization, providing epigenetic priming for subsequent immune sensing and effector transcriptional programs; ATAC-Fate2 was enriched in chromatin remodeling and gene expression–related epigenetic regulation, suggesting completion of landscape reconfiguration prior to transcriptional divergence and establishing a foundation for pathway-specific adjustments.

As differentiation progressed, the trajectories displayed pronounced functional polarization, with scATAC-seq and scRNA-seq results forming a coordinated “epigenetic regulation–transcriptional function” alignment that solidified path-specific polarization programs(Figure 4C, D).

In the late phase of RNA-Fate1, pathways such as MHC class II antigen processing and presentation, positive regulation of cytokine production, and inflammatory response regulation were markedly enriched, reflecting establishment of a core program centered on “antigen presentation–pro-inflammatory signal amplification–immune response enhancement. Correspondingly, ATAC-Fate1 showed enhanced chromatin accessibility in inflammation-associated cytokine production and positive regulation of the p38MAPK cascade, indicating systematic epigenetic activation of inflammatory signaling and cytokine programs that underpin stable transcriptional output. In contrast, late-stage RNA/ATAC-Fate2 was biased toward tissue homeostasis and immune modulation: RNA-Fate2 was enriched in efferocytosis (apoptotic cell clearance), positive regulation of cell adhesion, and immune effector process regulation—highlighting its role in microenvironmental homeostasis, immune modulation, and intercellular interactions. ATAC-Fate2 chromatin changes were strongly linked to positive regulation of cell differentiation, tissue development, and multicellular organismal development, establishing an epigenetic foundation for differentiation, maturation, and tis-sue-related functions.

Integrated multi-omics analysis demonstrated that monocyte differentiation into M1 pro-inflammatory and M2 regulatory macrophages follows a pattern of early epigenetic specialization followed by late functional polarization stabilization. Early in differentiation, both pathways shared transcriptional tendencies toward immune recruitment and initiation, while epigenetically initiating path-specific remodeling—ATAC-Fate1 focusing on immune perception–related regulatory element opening and ATAC-Fate2 emphasizing chromatin landscape reprogramming—creating coordinated early features of shared transcriptional priming and trajectory-specific epigenetic adaptation that set the stage for subsequent polarization. In the late phase, epigenetic features from scATAC-seq aligned directionally with scRNA-seq transcriptional output, consolidating into prototypical M1 polarization (“antigen presentation–proinflammatory amplification–immune enhancement”) and M2 polarization (“efferocytosis–tissue homeostasis–immune regulation”). This provides a clear epigenetic–transcriptional collaborative framework for the molecular trajectory of monocyte-to-specialized macrophage differentiation, offering precise multi-omics evidence for the mechanisms underlying macrophage functional heterogeneity in the ccRCC tumor microenvironment.

Employing the same strategy, we constructed early (C1-Net) and late (C2-Net) stage-specific transcriptional regulatory networks for each pathway, identifying core nodes whose functions aligned closely with the corresponding differentiation stage and delineating the complete regulatory chain (Figure 4E, F).

Along the monocyte-to-M1 macrophage path (RNA/ATAC-Fate1), NFIC constituted the central node in the early network (C1-Net). Overexpression of NFIC has been shown to promote monocyte differentiation and enhance cell survival^[32]^, consistent with the early immune response priming observed in RNA-Fate1 and laying the groundwork for subsequent M1 proinflammatory polarization. In the late network (C2-Net), IL1B—an inflammation-associated gene highly expressed in ccRCC M1 macrophages with anti-tumor potential^[33]^—emerged as the core hub, reinforcing the M1 proinflammatory immune response and solidifying the late pro-inflammatory phenotype.

Along the monocyte-to-M2 macrophage path (RNA/ATAC-Fate2), CEBPD served as the central node in the early network (C1-Net), appearing in both Fate1 and Fate2 early networks and suggesting bidirectional regulatory capacity. Existing studies indicate that CEBPD participates in both M1 and M2 polarization via the CE-BPD/RGS2/PAR2 axis^[34, 35]^, aligning with the shared early preparation and epigenetic specialization across pathways. In the late network (C2-Net), GLI2 acted as the key regulator of M2 polarization; its silencing reverses tumor-induced M2 polarization, while the SHH/GLI2-TGF-β1 loop promotes tumor progression^[36]^. This is consistent with the late-stage immune regulation and tissue homeostasis maintenance phenotype of Fate2, positioning GLI2 as the core hub for M2 phenotype consolidation.

Through integrated multi-omics analysis, we systematically delineate the dynamic trajectories and regulatory mechanisms governing monocyte differentiation into M1 pro-inflammatory and M2 immunosuppressive macrophages in the ccRCC tumor microenvironment. Both pathways share transcriptional programs for immune recruitment and effector priming early on, while epigenetic remodeling exhibits path-specific divergence, establishing the foundation for functional polarization. In the late phase, epigenetic regulation and transcriptional output achieve synergistic alignment, solidifying M1-type “antigen presentation–pro-inflammatory amplification–immune enhancement” and M2-type “efferocytosis–tissue homeostasis–immune regulation” phenotypes. Stage-specific regulatory networks further identify NFIC/IL1B and CE-BPD/GLI2 as pivotal core nodes, collectively forming a temporal regulatory axis of macrophage polarization and providing precise molecular evidence for the establishment of the immunosuppressive microenvironment in ccRCC.

### Result 5 Cell-Cell Communication Networks in ccRCC Progression Reveal Tumor Microenvironment Transition from Inflammation to Immunosuppression

To dissect the dynamic intercellular communication processes within the tumor microenvironment (TME) of clear cell renal cell carcinoma (ccRCC), we identified the 15 most active ligand-receptor interactions across different WHO grades using MultiNicheNet. By examining the grade-specific distribution of these core signaling axes, we delineated the evolutionary trajectory of the intercellular communication network(Figure 5A,B). However, given the inherent limitations of scRNA-seq in detecting low-abundance ligands or those derived from underrepresented cell populations, classical ligand-receptor analysis based solely on expression may underestimate function-ally relevant signaling pathways. To address this, we complemented the approach with a sender-agnostic ligand activity inference method, inferring ligand potential from the transcriptional response profiles of receiver cells across WHO grades(Figure 5C).

**Fig. 5.**
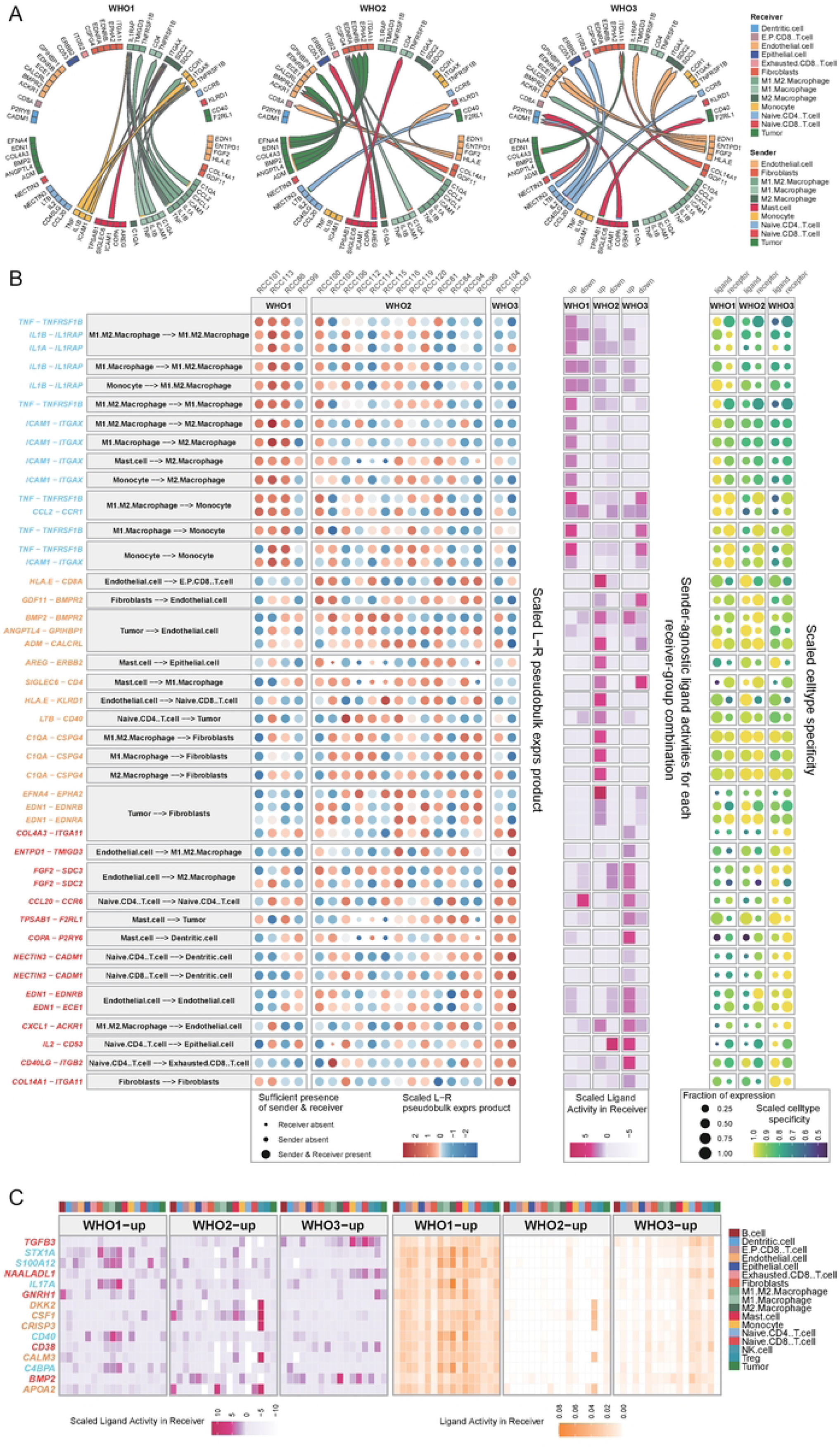
Stage-specific intercellular communication and ligand-receptor crosstalk in the ccRCC tumor microenvironment. A Circos plot showing the top 15 expressed ligand-receptor pairs across all cell types within the tumor microenvironment in different WHO grades, identifying the corresponding sender and receiver cell populations. B Combined bubble chart presenting the specific details of the top 15 interaction relationships across WHO classifications. The color of the interaction relationships indicates the specific classification they belong to. C Analysis of the top 5 sender-agnostic ligand activities across different tumor grades. **Alt text:** Diagrams and charts on intercellular communication in ccRCC, with subfigures labelled from A to C, illustrating ligand-receptor crosstalk via Circos plots, interaction bubble charts across WHO grades, and ligand activity analysis.

In WHO grade 1 tumors, intercellular communication was predominantly orchestrated by myeloid populations, including monocytes and macrophage subtypes. Top-ranked ligand-receptor pairs were enriched in classical pro-inflammatory pathways, such as IL1B–IL1RAP^[37]^, TNF–TNFRSF1B^[38]^, and ICAM1–ITGAX^[39]^, forming dense autocrine and paracrine loops among monocytes and macrophages. These interactions underscored a robust inflammatory TME in early ccRCC, characterized by immune activation and leukocyte recruitment. Consistent with this profile, sender-agnostic ligand activity analysis revealed elevated activity of inflammation-associated ligands—including IL17A^[40, 41]^, S100A12^[42]^, and CD40^[43, 44]^—in M2-like macrophages. Although these ligands were not among the most highly ex-pressed by sender cells in conventional ligand-receptor inference, their inferred activity based on receiver transcriptional responses confirmed persistent inflammatory pressure shaping macrophage functional states in early-stage disease.

In WHO grade 2 tumors, the communication landscape shifted markedly, with endothelial cells, fibroblasts, and tumor cells emerging as central signaling hubs. Dominant interactions included tumor-to-fibroblast EDN1–EDNRA/EDNRB^[45, 46]^, tumor-to-endothelial ADM–CALCRL^[47, 48]^, and matrix/vascular BMP2/GDF11–BMPR2 signaling^[49, 50]^, indicative of active vascular remodeling and stromal reprogramming during mid-stage progression.

Concurrently, sender-agnostic inference predicted heightened activity of immunosuppressive ligands in NK cells, notably DKK2, CSF1, and APOA2. DKK2 directly suppresses NK and CD8⁺ T cell activity to facilitate immune evasion^[51]^; CSF1 contributes to macrophage-mediated immunosuppression^[52]^; and APOA2 may promote broader immune tolerance via modulation of antigen presentation^[53, 54]^. These findings suggest incipient functional suppression of cytotoxic immune compartments at this transitional stage.

In WHO grade 3 tumors, the network was dominated by structural and homeo-static signals involving endothelial cells, fibroblasts, and tumor cells. Leading interactions encompassed extracellular matrix pathways such as COL14A1/COL4A3–ITGA11^[55, 56]^, endothelial autocrine EDN1–ECE1/EDNRB^[46, 57]^, and endotheli-al-to-M2 macrophage FGF2–SDC2/SDC3 signaling^[58, 59]^, reflecting aberrant vascular tone and matrix remodeling. Sender-agnostic analysis concurrently highlighted prominent immunosuppressive features.

Notably, elevated TGFB3 activity in naive CD8⁺ T cells implicated TGF-β signaling as a central suppressor of T cell function in advanced ccRCC^[59, 60]^. Heightened CD38-related signaling in Treg cells further reinforced an enhanced immunosuppressive T cell compartment in late-stage TME^[61, 62]^. In M2-like macrophages, increased BMP2 activity suggested contributions to immunosuppression and matrix remodeling via reinforced M2 polarization[63–65].

Collectively, these analyses delineate a stepwise TME evolution in ccRCC: from myeloid-driven inflammatory dominance in WHO grade 1, through emergence of vascular-matrix remodeling and incipient immune regulation in grade 2, to establishment of a stabilized immunosuppressive microenvironment in grade 3. Expression-based ligand-receptor inference delineates the structural framework of communication, while sender-agnostic ligand activity inference captures complementary functional signals manifested in receiver transcriptional programs.

### Result 6 Integrating machine learning algorithms to construct an optimal prognostic signature for ccRCC

Machine learning has demonstrated substantial promise in biomedical and oncologic research, particularly for tumor prognostication. By effectively capturing patterns in complex, high-dimensional data, machine learning can uncover prognostic signatures that conventional statistical approaches may fail to detect. In this study, we leveraged the Mime R package—which integrates 117 models derived from 10 distinct machine learning algorithms—to precisely identify prognostic features from clear cell renal cell carcinoma (ccRCC) tumor cells.

Building on prior findings, we next curated a biologically informed set of candidate genes for prognostic model construction. The top 15 regulatory nodes were selected from gene regulatory networks of tumor cells, T cells, and myeloid cells using maximum clique centrality (MCC). These nodes served as input features for subsequent machine learning analyses(Figure 6A).

**Fig. 6.**
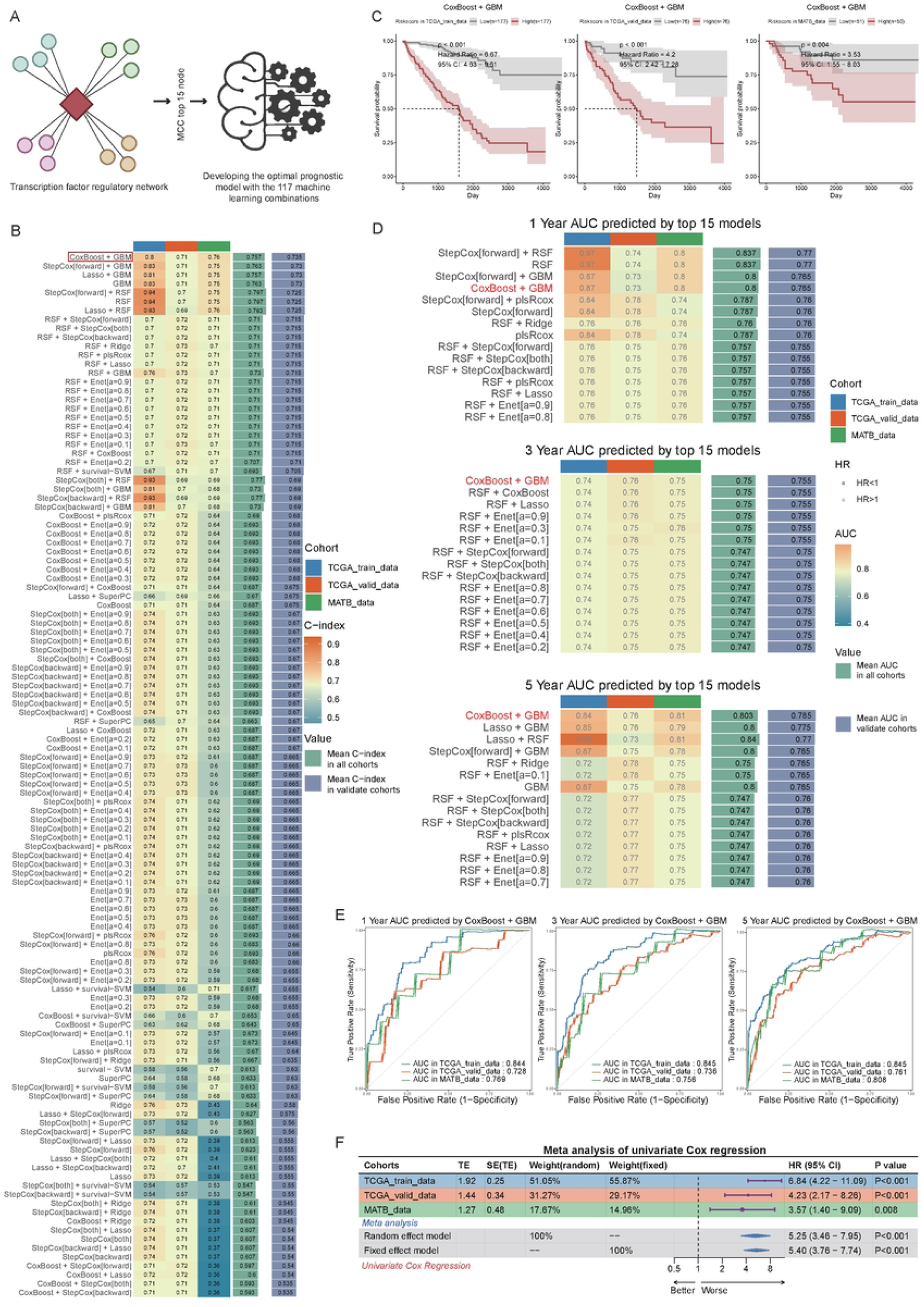
Construction of a robust prognostic signature based on ccRCC-associated molecular features. A Selection of the top 15 nodes in the transcription factor regulatory network based on MCC values for prognostic model construction. B C-index comparison of each candidate model across multiple datasets, sorted by the mean C-index of the validation cohorts. C Association between risk scores calculated by the CoxBoost + GBM combined model and patient prognosis in the training, validation, and additional validation cohorts. D Comparison of 1-year, 3-year, and 5-year AUC values for the CoxBoost + GBM combined model in different data sets. E Evaluation of 1-year, 3-year, and 5-year AUC performance consistency for the CoxBoost + GBM model across various datasets. F Meta-analysis of univariate Cox regression results for the CoxBoost + GBM combined model across all studied cohorts. **Alt text**: Graphs and data on the construction of a ccRCC prognostic model, with subfigures labelled from A to F, illustrating node selection, C-index performance, risk score survival associations, time-dependent AUC curves, and meta-analysis of model robustness.

The training cohort comprised survival data from 506 ccRCC patients in the TCGA database. We partitioned the dataset at a 7:3 ratio, randomly assigning 357 cases to the training set and 149 to the internal validation set. An independent external validation cohort (E-MTAB-1980; n = 101) was additionally included. Among the 117 models generated by Mime, the CoxBoost + GBM ensemble (CBG) achieved the highest mean concordance index (C-index) across the validation cohorts (TCGA validation and E-MTAB-1980), reflecting superior predictive performance (Figure 6B).

Using the median risk score derived from CBG, patients were stratified into high– and low-risk groups. Kaplan–Meier survival analysis demonstrated significantly poorer survival in the high-risk group compared with the low-risk group across all cohorts (Figure 6C). These findings indicate that the CBG model effectively discriminates patients with unfavorable versus favorable prognosis based on gene expression profiles alone.

To further assess model discrimination, we performed time-dependent receiver operating characteristic (ROC) curve analysis. CBG exhibited consistently high and stable area under the curve (AUC) values at 1, 3, and 5 years across the three cohorts (Figure 6D, E), underscoring robust generalizability.

Univariate Cox proportional hazards regression meta-analysis, implemented via Mime, confirmed that the CBG-derived risk score was a significant adverse prognostic factor in ccRCC (Figure 6F).

With the advent of next-generation sequencing, numerous machine learning-based prognostic signatures have been developed for ccRCC. To benchmark CBG against published models, we retrieved and evaluated multiple established signatures. Univariate Cox regression across all datasets revealed that CBG was most strongly associated with adverse outcomes in every cohort (Figure 7A). CBG also outperformed comparators in C-index across all datasets (Figure 7B) and ranked among the highest in 5-year AUC in nearly all cohorts (Figure 7C, Supplementary Figure S1H,I). Collectively, these comparisons highlight the superior extrapolation potential and bench-marking utility of CBG.

**Fig. 7.**
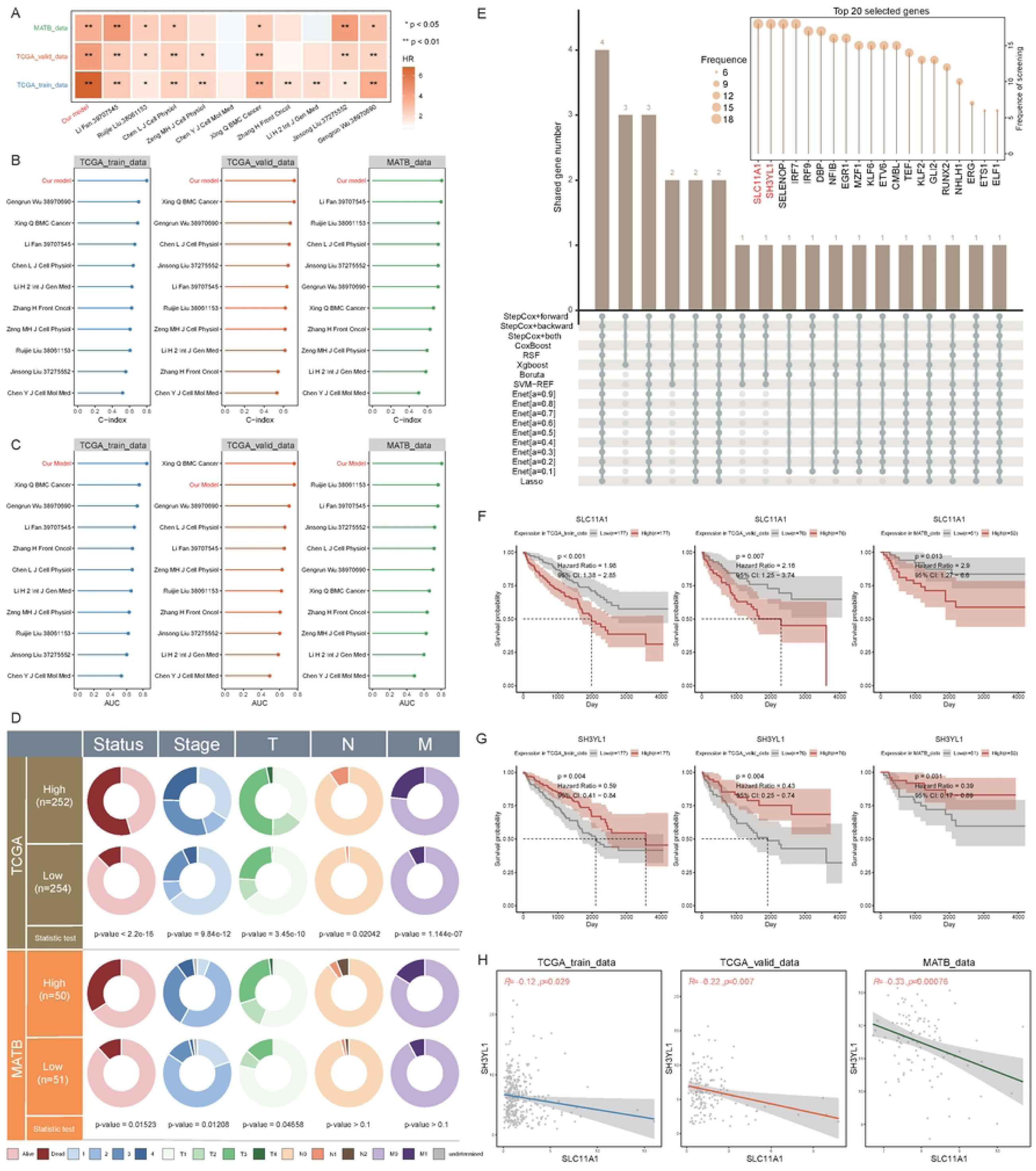
Performance evaluation and validation of the integrated CoxBoost and GBM machine learning model. A Comparison of hazard ratios (HR) between the CoxBoost + GBM model and 10 previously published models across three independent datasets. B Comparison of C-index performance between the CoxBoost + GBM model and 10 published models across three datasets. C Comparison of 5-year AUC values between the CoxBoost + GBM model and 10 published models across three datasets. D Correlation analysis between clinical characteristics and the high/low risk groups stratified by the CoxBoost + GBM model. E Core feature genes selected by various machine learning algorithms; the bottom histogram shows gene intersections across models, and the upper right panel displays gene selection frequency. F Kaplan–Meier overall survival curves of TCGA-KIRC patients stratified by the median expression of SLC11A1. G Kaplan–Meier overall survival curves of TCGA-KIRC patients stratified by the median expression of SH3YL1. H Pearson correlation analysis between SLC11A1 and SH3YL1 expression levels across different cohorts. **Alt text**: Graphs and data on the performance validation of the CoxBoost + GBM model, with subfigures labelled from A to H, illustrating benchmarking against published models (HR, C-index, AUC), clinical correlations, machine learning feature selection, and survival analysis of key genes SLC11A1 and SH3YL1.

Given that clinical parameters remain the mainstay of ccRCC prognostication in routine practice, we examined associations between CBG risk groups and clinicopathologic features. In the TCGA-KIRC cohort, high-risk patients exhibited significant positive correlations with higher WHO histological grade, overall clinical stage, and T, N, and M categories compared with low-risk patients (Figure 7D). In the E-MTAB-1980 validation set, high-risk status correlated significantly with grade, stage, and T category, although associations with N and M stages did not reach statistical significance—likely attributable to the smaller sample size and limited representation of advanced nodal (N1/N2, n = 7) and metastatic (M1, n = 12) cases, resulting in reduced statistical power.

To identify the most influential genes underpinning CBG performance, we conducted core feature selection using multiple algorithms (Figure 7E). SLC11A1 and SH3YL1 emerged as the genes most consistently selected across methods and were retained in our final model. We therefore focused on these two genes for deeper biologic interrogation.

Survival analysis revealed that high SLC11A1 expression was significantly associated with poorer overall survival across all three cohorts, whereas high SH3YL1 expression correlated with improved survival (Figure 7F, G). Prior studies have established SLC11A1 as a risk factor in ccRCC, with strong association to adverse prognosis in TCGA-KIRC^[66]^.

SH3YL1 has been implicated in renal injury pathways, particularly through interactions with NADPH oxidases such as NOX4^[67]^, and its downregulation is linked to worse overall survival in muscle-invasive bladder cancer. Notably, this study is the first to identify SH3YL1 as an independent prognostic factor in ccRCC, reinforcing its potential as a clinically relevant biomarker.

Pearson correlation analysis of tumor tissue expression profiles revealed a significant negative correlation between SLC11A1 and SH3YL1 (Figure 7H), suggesting that these genes may exert antagonistic effects during ccRCC tumorigenesis and progression.

In conclusion, we developed a robust prognostic model for ccRCC using a novel, biologically guided screening strategy. The resulting CBG signature effectively stratifies patients by survival outcome, exhibits strong discrimination across clinical features—including WHO grade—and demonstrates high stability and clinical relevance.

## Discussion

Clear cell renal cell carcinoma (ccRCC), the predominant subtype of kidney cancer, exhibits prognosis that is strongly influenced by WHO histological grade. Nevertheless, conventional grading systems based on morphology and bulk omics fail to adequately capture the multi-pathway malignant evolution of tumor cells, transcriptional regulatory heterogeneity, and the progressive remodeling of the tumor microenvironment (TME) from a pro-inflammatory to an immunosuppressive state. The present study employed an integrated single-cell multi-omics framework combining scRNA-seq and scATAC-seq to construct a comprehensive tumor ecosystem atlas spanning WHO grades 1 to 3 in ccRCC. This approach systematically revealed the malignant trajectories of tumor cells, CD8⁺ T cell exhaustion dynamics, bidirectional M1/M2 macrophage polarization, intercellular communication evolution, and the development of a prognostic model grounded in multi-perspective regulatory networks. Collectively, these findings provide high-resolution, dynamic molecular and immuno-logical evidence underpinning grade-specific progression in ccRCC.

A central discovery of this work is the multi-pathway evolutionary trajectory of tumor cells. We identified parallel differentiation routes—including RNA-Fate1/2 and ATAC-Fate1/3—wherein epigenetic alterations consistently preceded transcriptional reprogramming and functional adaptation. This “epigenetic pre-adaptation followed by transcriptional activation” paradigm aligns with emerging evidence across multiple cancer types, highlighting the anticipatory role of epigenetic regulation in malignant progression^[68]^. Stage-specific regulatory networks pinpointed RELA (early inflammatory driver), SLC6A3 (mid-stage HIF-associated marker), and ZBTB7A (late proto-oncogene) as pivotal nodes, each of which has been independently linked to adverse prognosis in ccRCC^[19, 22, 24]^. SLC6A3, a downstream effector of HIF signaling, occupied a central position in the mid-stage network, reflecting the hallmark metabolic reprogramming characteristic of ccRCC. In contrast, ZBTB7A dominated the late-stage network, consistent with its roles in promoting tumor invasion and survival. The sequential activation of these regulatory axes delineates a clear progression from early signaling sensitivity to late-stage metabolic maturity and enhanced invasiveness, offering a mechanistic framework for ccRCC heterogeneity.

At the immune microenvironment level, this study uncovered coordinated dynamics of CD8⁺ T cell exhaustion and macrophage polarization. The CD8⁺ T cell trajectory progressed from naive to intermediate exhaustion and subsequently to non-terminal exhaustion, with core regulatory nodes shifting from IRF7 (early activation and infiltration potential) to ZNF683 (mid-to-late plasticity maintenance)—a transition previously associated with responsiveness to PD-1 blockade^[26]^. The ZNF683-marked intermediate exhausted subset retains proliferative capacity, potentially representing a therapeutic window for immunotherapy^[69]^. Macrophage polarization followed dual paths driven by NFIC/IL1B (M1 pro-inflammatory) and CE-BPD/GLI2 (M2 immunoregulatory), with CEBPD implicated in tumor-induced bidirectional polarization^[34, 35]^; the present work extends this regulatory circuitry within the context of ccRCC grade progression. These immune cell trajectories interact synergistically with tumor cell evolution, collectively orchestrating the TME shift from early inflammation dominance to late immunosuppression.

Intercellular communication analysis reinforced this stepwise remodeling: WHO grade 1 featured myeloid-driven inflammatory signaling (e.g., IL1B/TNF/ICAM1 axes), grade 2 transitioned toward vascular-stromal remodeling (e.g., EDN1/ADM/BMP2 pathways), and grade 3 was dominated by immunosuppressive signals (e.g., TGFB3/CD38/DKK2). Sender-agnostic ligand activity inference uncovered latent inhibitory cues—such as DKK2-mediated suppression of NK/CD8 activity—that circumvented limitations inherent to classical ligand-receptor analysis^[70]^. These dynamic shifts align with prior reports of ccRCC TME evolution from proinflammatory to pro-tumorigenic states^[71]^, underscoring that grade progression arises not solely from intrinsic tumor cell changes but from holistic TME ecological adaptation.

The machine learning-based prognostic model (CBG), constructed from core nodes of multi-angle transcriptional regulatory networks, represents a key translational advance of this study. CBG demonstrated robust performance in both the TCGA training/validation sets and the independent E-MTAB-1980 cohort, outperforming existing signatures in C-index and time-dependent AUC. High expression of the core marker SLC11A1 correlated with poor prognosis^[66]^, whereas SH3YL1 showed the opposite association^[67]^. The significant negative correlation between these two genes suggests potential antagonistic mechanisms, opening new avenues for ccRCC biomarker research. Notably, CBG risk scores exhibited strong positive correlations with WHO grade and TNM stage, affirming clinical relevance.

Limitations of the study include the relatively modest sample size (19 pairs), which may constrain the full characterization of inter-grade heterogeneity^[12]^, as well as the absence of in vivo functional validation and multi-center cohort confirmation. Future investigations should expand to larger multi-omics cohorts and experimentally validate the therapeutic potential of key nodes (e.g., ZNF683, GLI2) through targeted interventions^[72]^.

## Data availability

All data used in this study are publicly available from GEO (https://ncbi.nlm.nih.gov/geo/) and TCGA (https://portal.gdc.cancer.gov/). The datasets analyzed are existing publicly accessible datasets. The single-cell RNA sequencing dataset used in this study is GSE207493. Furthermore, in addition to the data sets from the TCGA database, we also introduced an independent data set E-MATB-1980 (https://www.ebi.ac.uk/biostudies/arrayexpress/studies/E-MTAB-1980) consisting of 101 cohorts as an additional validation set for machine learning.

## Availability of Source Code and Requirements

Project name: ccRCC

Project homepage: https://github.com/Zhangyunpeng1987/ccRCC

Operating system(s): Platform independent

Programming language: R

Other requirements: R 4.3.0 or higher

License: MIT license

RRID: SCR_028348

## Additional Files

**Supplementary Fig. S1.**
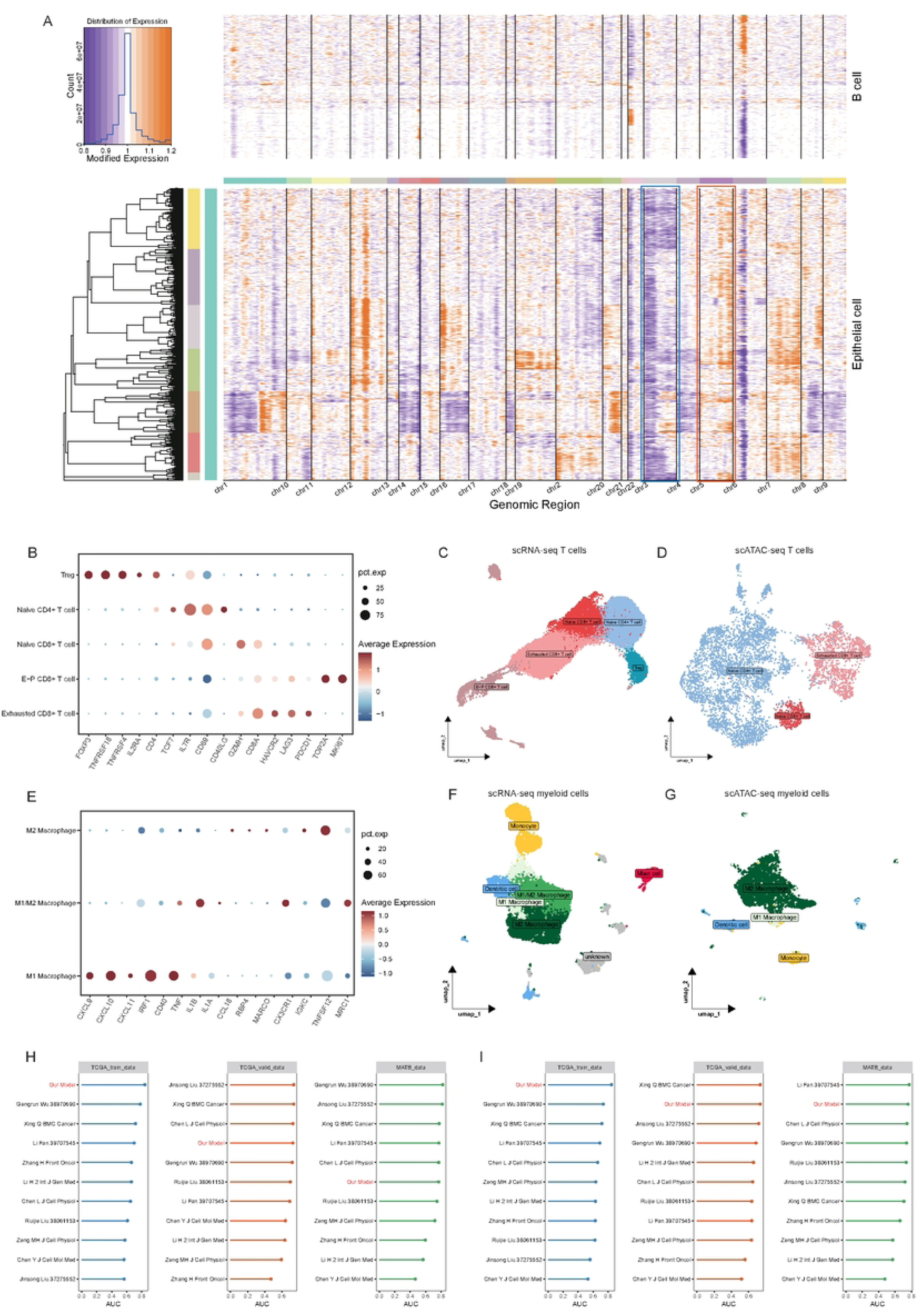
Characterization of copy number variations and cell-type annotation within the ccRCC landscape. (A) Copy number variation (CNV) analysis of epithelial cells in clear cell renal cell carcinoma samples, utilizing B cells as the reference population. (B) Dot plot showing the expression of canonical marker genes utilized for the systematic annotation of T cell subsets. (C, D) UMAP visualization of identified T cell subsets based on transcriptomic (scRNA-seq, C) and epigenetic (scATAC-seq, D) profiles. (E) Dot plot illustrating the expression of signature marker genes used for the identification of macrophage subsets. (F, G) UMAP projections showing the sub-clustering of myeloid cell subsets derived from scRNA-seq data (F) and scATAC-seq data (G). (H, I) Benchmarking of the prognostic performance (AUC) of the integrated CoxBoost + GBM model. The plots show a comparison of 1-year (H) and 3-year (I) AUC values between the proposed model and 10 previously published prognostic signatures across three independent datasets.

**Supplementary Table S1. Clinical profiles of the ccRCC patient cohort**. This table provides a comprehensive summary of the clinical and pathological characteristics for each patient with clear cell renal cell carcinoma (ccRCC) included in the current multi-omic study.

**Supplementary Table S2. Inventory of established prognostic models for ccRCC.** A detailed summary of previously published machine learning and statistical models for clear cell renal cell carcinoma prognosis, including their key features, algorithms, and validation metrics.

## CRediT authorship contribution statement

**Ruifei Liu:** Writing–review & editing, Validation, Methodology, Conceptualization. **Yuchen Shi:** Validation, Methodology, Investigation. **Yuxuan Xiao:** Writing–original draft, Methodology, Investigation. **Bolin Ren:** Validation; Investigation. **Liyuan Li:** Visualization, Investigation. **Bobin Qi:** Visualization; Investigation. **Tengyue Li:** Visualization; Investigation. **Yunpeng Zhang:** Conceptualization. **Jie Gao:** Conceptualization.

